# Field performance of the cardenolide-producing crucifer *Erysimum cheiranthoides* under herbivore attack and competition

**DOI:** 10.1101/2023.11.27.568809

**Authors:** Kunqi Wang, Erik van Bergen, Tobias Züst

**Affiliations:** Institute of Systematic and Evolutionary Botany, University of Zürich, Zollikerstrasse 107, CH-8008 Zürich, Switzerland; CE3C - Centre for Ecology, Evolution and Environmental Changes & CHANGE – Global Change and Sustainability Institute, Faculty of Sciences, University of Lisbon (FCUL), 1749-016 Lisbon, Portugal

**Keywords:** Enemy-free space, chemical novelty, plant defense, herbivore-imposed selection, plant-plant competition

## Abstract

Plants in the Brassicaceae produce glucosinolates as potent defenses against generalist herbivores, but many specialists can tolerate or deactivate these compounds. As a potential counter-adaptation, plants in the genus *Erysimum* evolved novel cardenolide defenses alongside ancestral glucosinolates. Here we grew the dually defended *E. cheiranthoides* and the phenologically similar *Rhamnospermum nigrum* in agricultural fields surrounded by Brassicaceae crops and monitored herbivore communities, leaf damage, and plant performance of focal plants over two seasons. In the second year, we additionally manipulated herbivore density and plant competition to identify direct and indirect effects on plant fitness. Both plant species were attacked by a diverse community of specialist herbivores, but most herbivores strongly preferred *R. nigrum*, reaching higher densities and causing intense damage that suppressed growth of these plants. A similar herbivore community attacked dually-defended *E. cheiranthoides*, but herbivores occurred at lower densities, caused less damage, and had no negative effect on growth. Consequently, herbivore suppression through pesticide application increased *R. nigrum* growth but provided no benefit to *E. cheiranthoides*. In fact, for *E. cheiranthoides* growing under heterospecific competition, pesticide application reduced growth due to increased competition by *R. nigrum*. Our results agree with a successful escape of *E. cheiranthoides* from herbivory, as even though plants are still attacked when herbivores are abundant, these herbivores caused little damage and caused no reduction in fitness. However, the escape from herbivory has likely come at loss of competitive ability when herbivore pressure is low, which would explain why despite its key innovation, *E. cheiranthoides* has been unable to gain dominance in any plant community.

## 1. Introduction

In nature, plants are constantly threatened by herbivores that consume a substantial proportion of global plant biomass and cause major yield losses in agriculture (Johnson 2011). To protect their limited resources, plants have evolved multiple layers of defense, including physical barriers and a vast array of chemical compounds (War et al. 2012).

Chemical defenses play a particularly important role in plant-herbivore interactions, as they shape herbivore community composition, and are themselves shaped by herbivore-imposed selection (Agrawal et al. 2012a, Züst et al. 2012). Over evolutionary time, reciprocal adaptations between plants and herbivores have produced a remarkable diversity of toxic, repellent, or antinutritive compounds across plant lineages (Speed et al. 2015).

Defense traits require substantial energetic and metabolic investment; for example, glucosinolate production in Brassicaceae species can consume up to 15% of a plant’s total photosynthetic energy (Bekaert et al. 2012). Defense traits therefore represent major costs for plants, both directly through reduced allocation to other functions (direct or allocation costs, Herms and Mattson 1992, Züst and Agrawal 2017) and indirectly by altering interactions with mutualists or competitors (ecological costs, Strauss et al. 2002). However, both costs and benefits of defense depend on the environmental context. For example, the growth-differentiation balance (GDH) hypothesis predicts a physiological trade-off between growth and defense, but rapid growth should be favored particularly during plant establishment (Herms and Mattson 1992, Züst et al. 2011), and investment in defense should be favored only when herbivory is intense and when herbivores are susceptible to that defense (Simms and Rausher 1987).

The interplay between competition, defense, and herbivory has been repeatedly demonstrated in experimental systems (Hambäck and Beckerman 2003). Manipulating herbivore pressure often leads to striking shifts in defense expression and plant-plant competition (Mauricio and Rausher 1997, Züst et al. 2012). The natural gains of novel defenses can mirror the effects of experimental herbivore exclusion: the acquisition of a potent new defense may allow plants to temporarily escape herbivore-imposed selection, thereby altering competitive dynamics and facilitating colonization of new habitats (“escape and radiate”, Ehrlich and Raven 1964).

In the Brassicales, the evolution of glucosinolate defenses approximately 80 million years ago triggered rapid species diversification (Edger et al. 2015). Over evolutionary time, specialist herbivores evolved counter-adaptations that allow them to tolerate or even exploit these defenses (Ratzka et al. 2002, Wittstock et al. 2004, Sporer et al. 2021). Most extant species of the Brassicaceae are therefore attacked by diverse communities of glucosinolate-resistant herbivores (Bidart-Bouzat and Kliebenstein 2008, Mertens et al. 2021). Yet despite limited effectiveness against specialists, Brassicaceae still produce a diverse array of glucosinolates (Fahey et al. 2001), highlighting the challenges of inferring early selective pressures that shaped defense evolution from present-day interactions.

Because long-term coevolution can obscure the processes that initially selected for new resistance traits, focusing on defenses of recent evolutionary origin provides a valuable opportunity to study costs and benefits before specialized antagonists evolve. In one such example, species of the genus *Erysimum* (Brassicaceae) evolved the ability to synthesize cardenolide toxins within the last five million years (Züst et al. 2020). Cardenolides inhibit animal Na^+^/K^+^-ATPases and therefore differ fundamentally from glucosinolates (Agrawal et al. 2012b). Consequently, some glucosinolate-resistant herbivores are completely repelled by *Erysimum* plants (Sachdev-Gupta et al. 1993, Younkin et al. 2024, but see Wang and Züst 2025). This apparent ‘escape’ from herbivores may have triggered a recent radiation within this diverse genus (Züst et al. 2020). Most *Erysimum* species co-express glucosinolate and cardenolide defenses, but while interspecific variation in glucosinolates is limited, cardenolide profiles are highly variable (Züst et al. 2020), potentially reflecting differential selection on the novel defense. To date, no herbivore has evolved specialized resistance to *Erysimum* cardenolides, and only one cardenolide-specialized seed bug has expanded its host range to include *Erysimum* (Petschenka et al. 2022). This system therefore offers a rare opportunity to study the ecological consequences of a newly evolved defense before coevolution could reshape it.

Here, we use the annual *Erysimum cheiranthoides* to characterize the natural herbivore community attacking this species in central Europe. Specifically, we ask whether the evolution of cardenolide production has allowed *E. cheiranthoides* to escape from herbivory, and aim to identify the selective forces acting on this species in its putative ‘enemy-free space’ (Jeffries and Lawton 1984). To quantify herbivory experienced by typical Brassicaceae species under these same field conditions, we use the annual *Rhamnospermum nigrum* (syn. *Brassica nigra*) as a phytometer (Clements and Goldsmith 1924). Phytometers are standardized transplanted species used to compare environments, and can be used to assess herbivore pressure across sites and years (Scherber et al. 2006). *R. nigrum* is readily attacked by glucosinolate-resistant herbivores, and shares a similar phenology, growth rate, and size with *E. cheiranthoides* under field conditions. Although the two species are phylogenetically distant, *R. nigrum* may thus functionally resemble a cardenolide-free ancestor of *E. cheiranthoides*.

Plants were grown in fields adjacent to oilseed rape (*Brassica napus*), which provided a high density of Brassicaceae-associated herbivores. We monitored herbivore abundance and plant damage across full growing seasons in two years. In the second year, we additionally applied insecticide treatments to manipulate herbivore pressure, and we included a competition treatment to assess how herbivory and defense interact to influence plant performance. We hypothesize that the novel cardenolide defense renders *E. cheiranthoides* better protected and less damaged by herbivores than *R. nigrum*, resulting in higher fitness. Conversely, we predict that under low herbivore pressure, the dual investment in defense may make *E. cheiranthoides* competitively inferior to *R. nigrum*.

## 2. Methods

### 2.1 Plant material

*E. cheiranthoides* is an annual plant native to central and northern Europe and parts of Asia (Polatschek 2010, 2013). Flowering occurs from May through September, with peak flowering in June (Polatschek 2013; T. Züst, personal observation). We grew a set of natural accessions of *E. cheiranthoides* (‘genotypes’ hereafter) from locations in western and central Europe in common garden experiments in Switzerland in 2022 and 2023. A total of 38 genotypes were used in 2022, but due to large phenological variation, a subset of nine late-flowering genotypes was selected for 2023. All genotypes were initially propagated for two generations under common greenhouse conditions. Seeds of *R. nigrum* used in this study were originally provided by the Botanical Garden of Krefeld, Germany.

Plants for field experiments were sown in late April for transplantation to the field in late May. Seeds of *E. cheiranthoides* were soaked and cold-stratified for three days at 4 °C, and three seeds were placed on the soil surface of pots (7 × 7 × 8 cm) filled with a 1:1 sand-peat substrate (Einheitserde, Patzer Erden GmbH, Germany). Seeds of *R. nigrum* were sown directly into pots and lightly covered with soil. Pots were maintained for germination under high humidity in a growth chamber for one week (constant light, 23 °C, 60% relative humidity), then excess seedlings were removed, and pots were transferred to an open greenhouse for hardening prior to field transplantation. In the last week of May, we transplanted three-week-old plants into freshly tilled fields at the Swiss federal agricultural research station (Agroscope Reckenholz, Switzerland). Plants were transplanted to the field inside their plastic pot to restrict root expansion and limit biomass growth. A small set of plants was kept in the greenhouse for an additional week and used for the assessment of leaf traits (trichome density and trichome size) and innate herbivore preference under laboratory conditions (see Supplementary Methods).

### 2.2 Field experiments

#### 2.2.1 Field experiment 2022

In 2022, we used a 6 × 80 m field site (Figure S1a, 47.4376° N, 8.5268° E) adjacent to a field of oilseed rape. On either side of the experimental plot, 1 m-wide buffer strips of a commercial cover crop mixture consisting of *Raphanus sativus* and *Avena strigosa* had been sown two months earlier. We divided the experimental plot into ten blocks, and in each block, we planted 38 *E. cheiranthoides* and four *R. nigrum* plants in a randomized grid pattern, with plants spaced 1 m apart and with a 0.5 m distance to the plot edge on each side. Throughout the experiment, plants were watered as needed, and the surrounding soil was weeded regularly and kept free of other vegetation. Unfortunately, buffer strips had to be mowed halfway through the experiment (week 5) to prevent weed seed dispersal.

In weeks 3, 6, and 9 after transplantation, we assessed plant damage by herbivores, and in the six interspersed weeks, we recorded the herbivores that were present on each experimental plant. During herbivore assessments, we surveyed each plant and recorded all herbivores by species or functional group. Herbivore assessments were completed over 2–3 consecutive days in each week. During damage assessments, we surveyed each plant to record leaf damage and plant size. Damage was quantified as the number of leaves with visible herbivory, and plant size was quantified as the total number of leaves and as plant height. A representative sample of leaves was harvested and analyzed to determine nutritional quality of plants throughout the 2022 experiment (see Supplementary Methods).

Whole plants were sequentially harvested for biomass, with complete blocks assigned to different harvests. Plants in blocks 1-3, 4-6, and 8-10 were harvested in weeks 8, 11, and 14, respectively. Aboveground biomass was cut at soil level, dried (60 °C) for 48–96 hours, and weighed to the nearest mg. Plants in block 7 were removed from the field in week 8 to complete seed maturation in a greenhouse. Seeds and vegetative biomass were dried and weighed separately, and mean seed mass was estimated from 100 seeds per plant.

#### 2.2.2 Field experiment 2023

We set up a modified field experiment in 2023 with added treatments of plant competition and herbivore suppression. The competition treatment consisted of *E. cheiranthoides* grown together with *R. nigrum* in the same pot, while the herbivore suppression treatment consisted of weekly applications of lambda-cyhalothrin (Kendo® Gold, Maag Gardens, Switzerland), a non-systemic pyrethroid insecticide.

We used a different 4 × 66 m field site (Figure S1b, 47.4284° N, 8.5232° E), again adjacent to a field of oilseed rape. On either side of the experimental plot, 1.5 m-wide buffer strips of the same cover crop mixture had been sown one week prior to transplanting, which were maintained throughout the experiment. We divided the experimental plot into six blocks, and for blocks 1, 2, 5, and 6, we planted 36 individual *E. cheiranthoides* and 4 *R. nigrum* plants each in a stratified random grid pattern as in 2022. For blocks 3 and 4, we planted 40 paired *E. cheiranthoides* and *R. nigrum* each in the same grid pattern (Figure S1b). Lambda-cyhalothrin (15 mg/L) was applied weekly for seven weeks to plants in blocks 2, 4, and 6, increasing spray volume with plant size (30-150 mL per plant). Plants in blocks 1, 3, and 5 were sprayed with an equivalent amount of water.

Herbivore assessments were carried out on individual plants as before (weeks 1, 2, 4, 5, 7, 8), while competing plants could only be assessed in the first half of the experiment.

Damage was assessed on all plants (individual plants: weeks 3, 6, 9; competing plants: weeks 3, 6) using a modified approach to distinguish leaves with high (leaves with more than 75% of area damaged), low, and no damage. Plant height was measured weekly, while during damage assessments, we also measured plant diameter and branch number. Because total leaf counts showed large observer bias in 2022, leaf numbers in weeks 6 and 9 were estimated in 2023 using predictive models based on 20 randomly sampled plants (see Supplementary Methods). All plants were harvested in week 10 by cutting the stems at soil level and placing the entire aboveground biomass in paper bags, followed by drying and weighing.

### 2.3 Statistical analyses

The current study focuses on species differences between *E. cheiranthoides* and *R. nigrum*, while accounting for variation among plant genotypes by including genotype as a random effect in all analyses. Genotype-level variance was quantified using intraclass correlation coefficients (ICC, function *icc*, R package ‘performance’; Lüdecke et al. 2021) for each trait. To ensure comparability between years, we only used data for *E. cheiranthoides* from the nine genotypes grown in both years of field experiments (and all data for *R. nigrum*). All analyses were performed in the statistical software R v.4.5.1 (R Core Team 2023). Mixed models were fitted using lme4 (functions *lmer*, *glmer*; Bates et al. 2015), generalized additive models (GAMs) using mgcv (function *gam*; Wood et al. 2016), and post hoc comparisons were performed using emmeans (function *emmeans*; Lenth 2023). Significance of fixed effects in mixed models was assessed using Type II or Type III Wald Χ^2^ tests (function *Anova* in R package ‘car’; Fox and Weisberg 2019).

#### 2.3.1 Herbivore community composition and phenology

To compare herbivore communities between plant species, we first analyzed total herbivore abundance (summed within year) and compared community composition at peak abundance using multivariate analyses (see Supplementary Methods). To examine temporal variation in herbivore presence and abundance, we then used a series of GAMs. Given the differences in experimental design between 2022 and 2023, we performed separate GAMs for each year. For each of the eight major herbivore species or species groups, we fitted two sets of GAMs. First, we fitted binomial GAMs to presence-absence data across all six weeks of observations, and second, we excluded weeks with zero observations for each herbivore and fitted negative binomial GAMs to abundance data of the remaining weeks. Nested GAMs differing in smoothing structure were fitted with maximum likelihood (ML), models were compared using AIC, and the selected models were refitted using REML (see Supplementary Methods). For 2022, GAMs included plant species as a fixed effect, whereas for 2023, we additionally included insecticide treatment and its interaction with plant species. Due to incomplete sampling of plants in the competition treatment, they were excluded from this analysis.

#### 2.3.2 Plant damage assessments

Leaf damage was analyzed using binomial generalized linear mixed-effects models fitted to the proportion of damaged leaves. For 2022, damaged and undamaged leaf counts per plant were used as a two-vector response representing the proportion of damaged leaves. Models included fixed effects of week, plant species, and their interaction, observer as a covariate, and random intercepts for plant identity and plant genotype.

For the 2023 field experiment we used a similar approach, but because damage levels were overall higher in this experiment, we used a two-vector response of highly damaged leaves (defined as > 75% leaf area damaged) *vs.* all undamaged and lightly damaged leaves.

This response was modeled using fixed effects of assessment week, plant species, insecticide treatment, and competition treatment, including all possible interactions. Because damage was not assessed for competing plants in week 9, we excluded the four-way interaction and specified competition interactions only with plant species and insecticide treatment. We again specified observer as a covariate, and included random intercepts for plant identity, neighborhood group (grouping together paired plants of the competition treatment), and plant genotype.

#### 2.3.3 Effects of herbivory and competition on plant growth

Herbivore numbers dropped substantially following the early mowing of buffer strips in 2022. We tested whether this reduction influenced plant growth using repeated height measurements taken in weeks 3, 6, and 9. For 2023, we used a similar analysis of weekly height measurements to assess the effect of insecticide application on growth (see Supplementary Methods).

For the 2022 field experiment, we analyzed the biomass of plants harvested in weeks 8, 11, and 14 using a linear mixed-effects model. Ln-transformed dry biomass was modeled using fixed effects of plant species, harvest week, and their interaction, and a random intercept of plant genotype. For the 2023 field experiment, we modeled ln-transformed biomass of plants (all harvested after 10 weeks) using plant species, insecticide treatment, competition treatment, and their interactions as fixed effects, with random intercepts for plant genotype and neighborhood group.

## 3. Results

### 3.1 Herbivore community composition and phenology

Across two years, we recorded 11 herbivore species or groups. All herbivores were found on both plants in both years, except for a few low-abundance species missing from either *R. nigrum* or *E. cheiranthoides* in 2023, indicating a largely consistent community within the oilseed rape–dominated landscape. Most herbivores were more abundant on *R. nigrum* than on *E. cheiranthoides* (Tables S1–S2). In 2023, insecticide treatment did not uniformly reduce herbivores: flea beetles and shield bugs decreased on *E. cheiranthoides*, whereas pollen beetles, whiteflies, and weevils increased on *R. nigrum*, and aphids increased on both species (Table S2).

Despite the shared herbivore community, community composition (estimated using Bray-Curtis dissimilarities) differed substantially across plant species, treatments, and years (see Supplementary Results, Figure S2). GAMs revealed pronounced seasonal dynamics (Figure 1), with aphids and flea beetles colonizing experimental plants first and reaching higher abundance on *R. nigrum*. Pollen beetles reached peak abundance by week 4, and most other herbivores by week 5. In 2022, six of eight major herbivores were more abundant on *R. nigrum*; four showed the same pattern in 2023. The only exceptions were crucifer shield bugs and cabbage whiteflies, which occurred at similar frequencies on both plant species in both years. Effects of *E. cheiranthoides* genotype on temporal herbivore dynamics were mostly modest (Table S4). In 2022, most herbivores declined sharply after week 5, a pattern not observed in 2023 (Figure 1). Additionally, GAMs revealed that herbivores on insecticide-treated plants not only reached higher peak abundances, but also persisted for longer on these plants (Figure S4).

**Figure 1.**
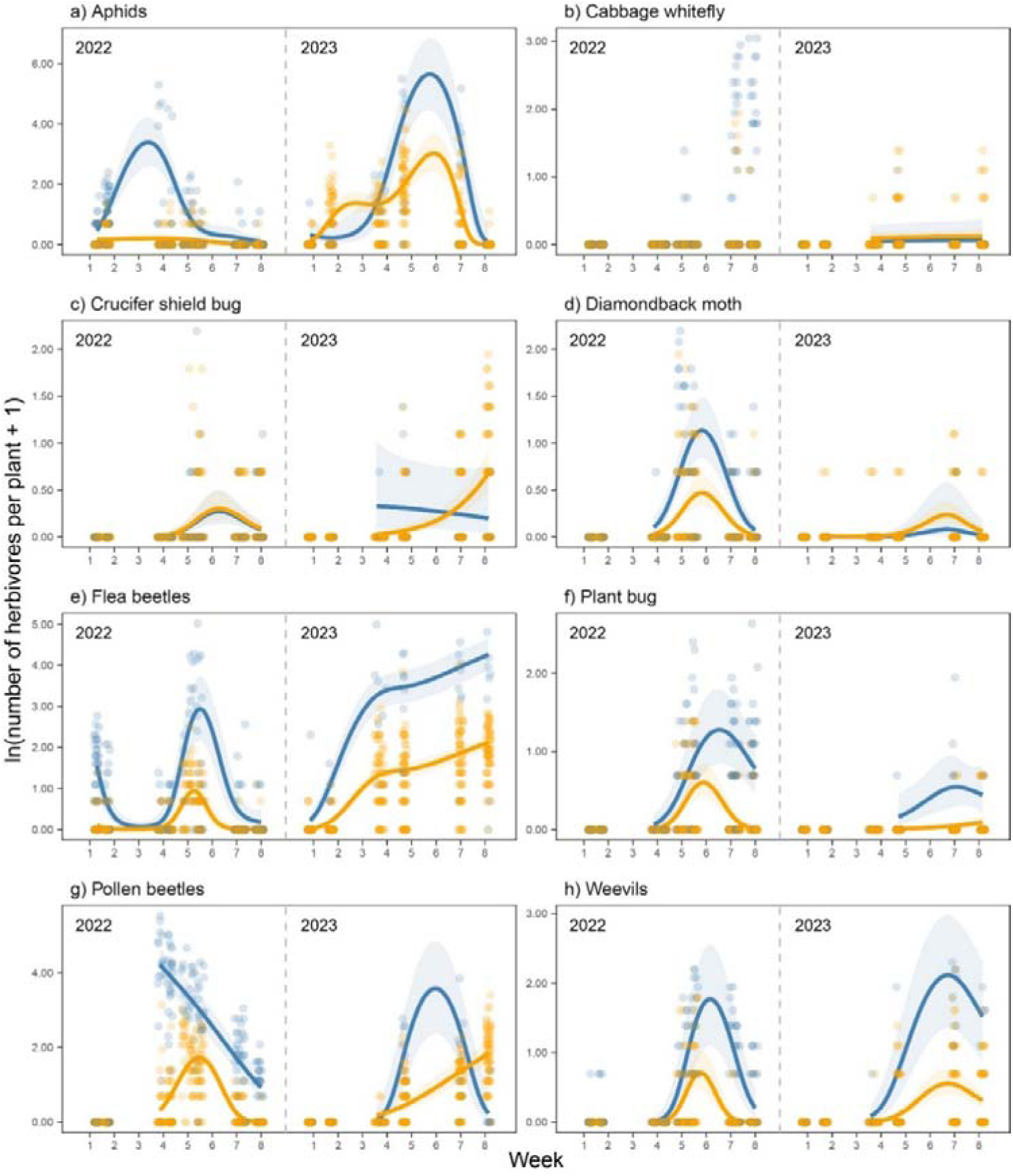
a-h) Herbivore abundance for the eight main herbivores observed on *R. nigrum* (blue) and *E. cheiranthoides* (orange) in the 2022 and 2023 field experiments. Points represent observations on individual plants, while lines and shaded areas are mean model predictions and 95% confidence intervals from negative binomial generalized additive models (GAMs). GAMs were fitted to count data after exclusion of weeks with zero observations, with separate models fitted for each year. No model could be fitted for cabbage whiteflies in 2022 due to insufficient observations on *E. cheiranthoides*. See Table S4 for test statistics of each model.

### 3.2 Plant damage

In 2022, leaf damage was substantially higher on *R. nigrum* than on *E. cheiranthoides* (Figure 2a; Table S5a), with only a modest proportion of variance attributable to plant genotype (ICC = 6.7 %). Damage to *R. nigrum* was nearly complete in weeks 3 and 6, whereas less than 30% of leaves were damaged on *E. cheiranthoides*. Damage declined on both species by week 9 (week 6 vs. 9: z-ratio = 13.17, p < 0.001), mirroring declining herbivore abundances.

**Figure 2.**
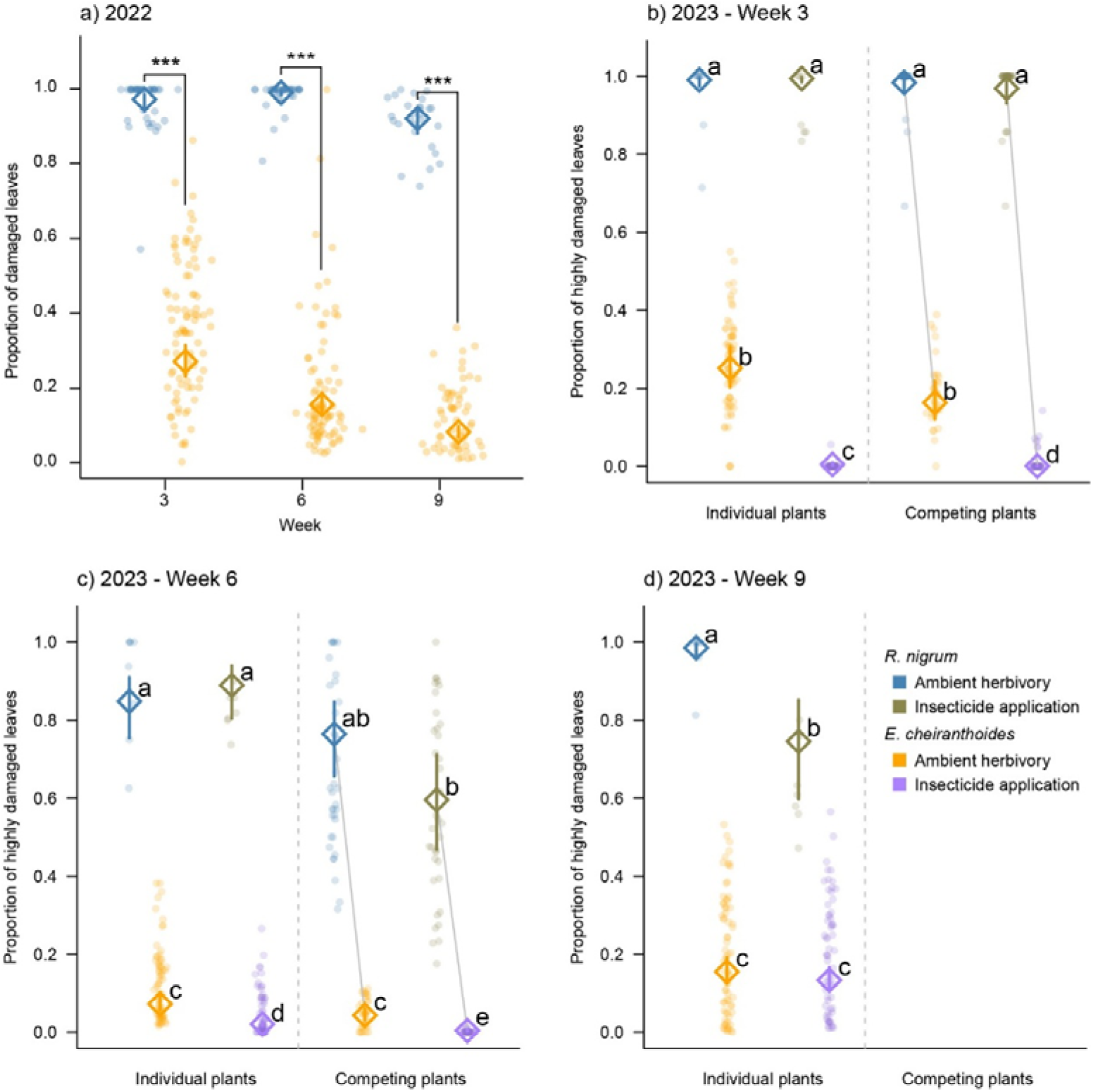
a) Proportion of damaged leaves in the 2022 field experiment. Points are values for individual plants, while diamonds and vertical lines are the means and 95% confidence intervals from a binomial GLMM. Brackets highlight significant linear contrasts within each week at the p < 0.05 level. b-d) Proportion of highly damaged leaves (leaves with more than 75% area damaged) on plants in weeks 3, 6, and 9 of the 2023 field experiment. Letters highlight significant linear contrasts at the p < 0.05 level within each week. Means of plants in the competition treatment are connected by grey lines to highlight the paired design. In week 9, damage was only quantified on individual plants.

In 2023, the proportion of heavily damaged leaves was higher on *R. nigrum* than on *E. cheiranthoides* (Figure 2b-d), with effects varying by treatment and week (Table S5b, plant genotype ICC = 4.3 %). Under ambient herbivory, damage levels remained approximately constant throughout the season. Insecticide treatment strongly reduced damage to *E. cheiranthoides* early in the experiment, whereas effects of insecticide on *R. nigrum* were only evident late in the season. While plant-plant competition did not significantly affect damage under ambient herbivory, the competition treatment amplified the effect of the insecticide treatment and reduced damage levels below those of the insecticide treatment alone in weeks 3 for *E. cheiranthoides*, and in week 6 for both plant species (Figure 2b-c). Week 9 lacked damage quantifications on competing plants; thus, competition contrasts could not be estimated for the final observation.

### 3.3 Plant growth and competition

In 2022, *R. nigrum* showed minimal height growth in the first half of the experiment during peak herbivore abundance (Figure S5a; weeks 3-6: absolute growth rate: 0.32 [0.30 – 0.34] cm day^-1^; mean [95% confidence interval]), whereas *E. cheiranthoides* grew more than five times faster in the same period (1.71 [1.62 – 1.79] cm day^-1^). Growth of *R. nigrum* increased only towards the end of the experiment, and between weeks 6 and 9, both species exhibited similar growth rates (Figure S5a, Table S6a). In 2023, both species grew faster under the insecticide treatment (Table S6b), with sprayed *R. nigrum* showing more than twofold higher growth between weeks 6 and 9 compared to controls.

In 2022, species differences in biomass shifted between early and later harvests (Table S7a), with a considerable proportion of variance attributable to plant genotype effects (ICC = 24.8 %). At the first harvest (week 8), *R. nigrum* had significantly lower biomass than *E. cheiranthoides* (Figure 3a, t-ratio = −3.02, p = 0.017). This difference disappeared by weeks 11 and 14, consistent with the increased late-season growth of *R. nigrum*. In 2023, biomass showed a significant species × insecticide × competition interaction (Figure 3b; Table S7b, plant genotype ICC = 8.4 %). Under ambient herbivory, biomass of *R. nigrum* was low, while *E. cheiranthoides* was largely unaffected. Biomass of *R. nigrum* was further reduced by competition under ambient herbivory but increased by the insecticide treatment irrespective of competition. In contrast, biomass of *E. cheiranthoides* was unaffected by competition or insecticide treatment individually, but reduced by their combined effects. Finally, aboveground biomass was strongly correlated with seed production in *E. cheiranthoides* in 2022 (Figure S6), suggesting biomass differences likely translate into fitness differences.

**Figure 3.**
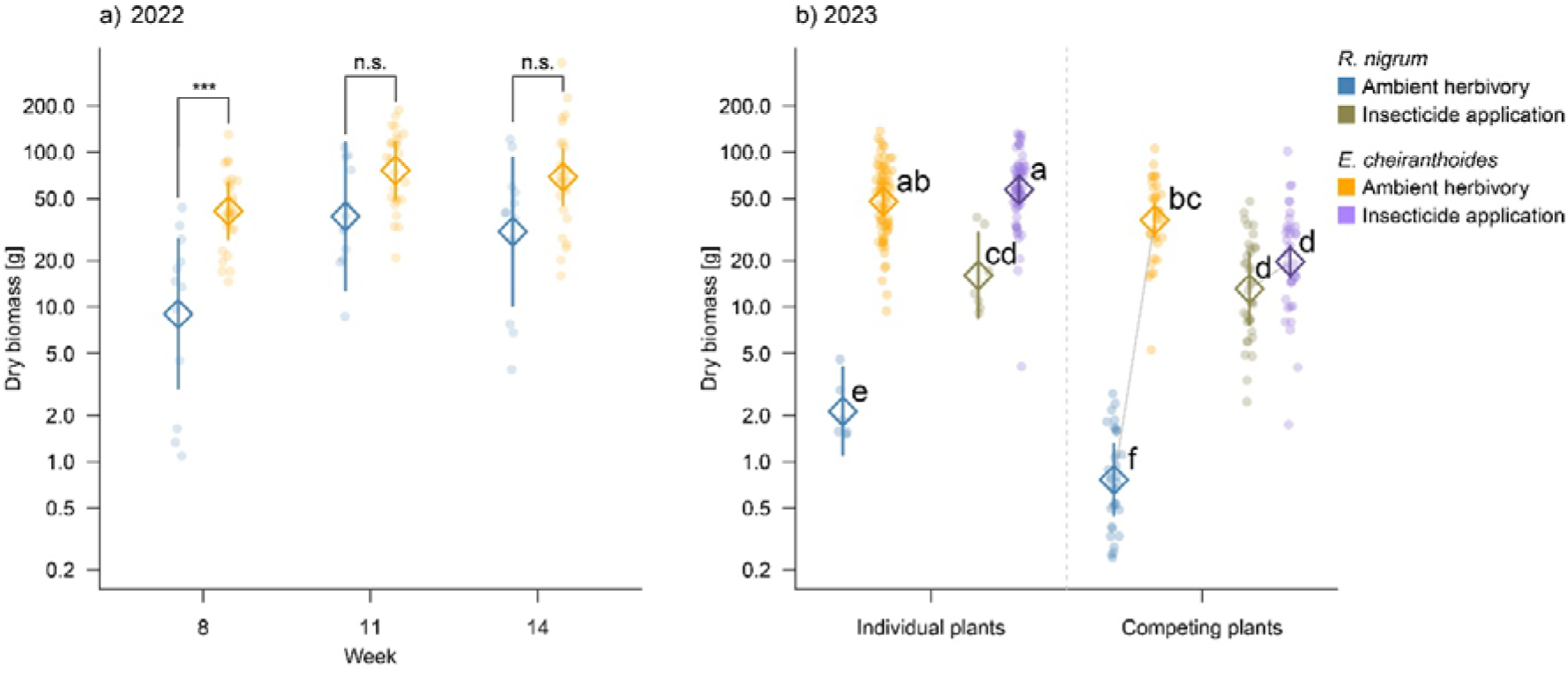
a) Total aboveground biomass of subsets of plants harvested after 8, 11, and 14 weeks in the 2022 field experiment. Diamonds and lines are means and 95% confidence intervals from a linear model. Brackets highlight significant linear contrasts within each week at the p < 0.05 level. b) Biomass of individual *R. nigrum* and *E. cheiranthoides* plants or heterospecific pairs harvested after 10 weeks of growth in the 2023 field experiment. Points are individual plant values, while diamond symbols and lines are means and 95% confidence intervals from a linear mixed-effects model. Letters denote significant linear contrasts at the p<0.05 level.

### 3.4 Plant traits and innate herbivore preference

Nutritional traits differed modestly between species (see Supplementary Results, Figure S7). In contrast, trichome density and size varied substantially, with leaves of *E. cheiranthoides* densely covered in small trichomes on both sides, while *R. nigrum* primarily expressed larger trichomes along leaf veins on the underside of its leaves (see Supplementary Results, Figure S8). In leaf-disc assays, all four tested herbivores preferred *R. nigrum* over *E. cheiranthoides* (Figure S9). This preference was least pronounced for *E. ornata*, mirroring the weaker preference for *R. nigrum* of this herbivore in the field.

## 4. Discussion

*E. cheiranthoides* experienced substantially less herbivory than *R. nigrum* across two field seasons. While severe herbivore damage suppressed growth in *R. nigrum*, growth of *E. cheiranthoides* remained largely unaffected even at peak herbivore abundance, consistent with escape from strong herbivore-imposed selection. However, manipulating herbivore abundance and competition revealed that *E. cheiranthoides* was competitively suppressed by *R. nigrum* when herbivory was reduced, indicating that the defensive superiority of *E. cheiranthoides* is coupled with relatively weak competitive ability.

Herbivores observed on both plant species were mostly glucosinolate-resistant Brassicaceae specialists, likely originating from surrounding crops and buffer strips and spilling over onto our plots (associational susceptibility; Barbosa et al. 2009). This landscape effect is illustrated by the sharp decline in herbivores following the premature mowing of buffer strips in 2022. Although we selected a Brassicaceae-dominated environment to ensure high specialist herbivore pressure, such dominance is common in Brassicaceae field studies (Bidart-Bouzat and Kliebenstein 2008, Mertens et al. 2021), likely because glucosinolates filter out many generalist herbivores (but see Zalucki et al. 2021).

We never recorded larvae or eggs of the cabbage white butterfly (*Pieris rapae*), despite frequent adult observations at the field sites. Absence on *E. cheiranthoides* is consistent with their aversion to cardenolides (Sachdev-Gupta et al. 1993), but absence on *R. nigrum* was unexpected. One possibility is that *R. nigrum* gained field-scale associational resistance from growing in proximity to *E. cheiranthoides* (Atsatt and O’Dowd 1976, Barbosa et al. 2009), which is further supported by the observed reduction in damage on insecticide-treated *R. nigrum* paired with *E. cheiranthoides*.

Most herbivores were more abundant on *R. nigrum* than on *E. cheiranthoides* across feeding guild. An exception were crucifer shield bugs (*Eurydema spp.*), which occurred at similar abundances on both species. Although *Eurydema* commonly feed on *Erysimum* species (Gómez 2005), our laboratory choice assays still showed a slight preference for *R. nigrum*, and *Eurydema* remain broadly generalized across the Brassicaeae (Piersanti et al. 2020). Whether *Eurydema* actively prefers *Erysimum* under natural conditions thus remains unresolved.

None of the observed herbivores completely avoided *E. cheiranthoides*, but importantly, we did not distinguish between perching and feeding. Highly mobile taxa (e.g., pollen beetles, flea beetles) may frequently alight but depart after brief sampling. Such mobility may also explain higher herbivore counts on insecticide-treated plants in 2023, as weekly applications likely reduced residents while increasing attractiveness to transient visitors. Consistent with an overall positive insecticide effect, sprayed plants were less damaged in some weeks and reached higher biomass by the end of the experiment.

Although our study compares only two species, we included *R. nigrum* as a phytometer – a standardized, glucosinolate-defended Brassicaceae used to estimate background herbivore pressures and competitive context. Because it is broadly attractive to glucosinolate-resistant specialists and matches *E. cheiranthoides* in phenology and size, *R. nigrum* provides a practical benchmark for ‘cardenolide-free’ Brassicaceae performance.

Nonetheless, because many traits differ between species, we cannot attribute reduced herbivory solely to cardenolides. For example, while nutritional differences between the species were modest, trichome density and structure differed markedly. Trichomes often play a role in plant defense against insects (Levin 1973, Sato et al. 2019), and may be particularly important for the *Erysimum* system (Wang et al. 2025). Even though cardenolides remain a key distinguishing trait between the two species, additional roles for trichomes and other unmeasured traits cannot be excluded.

A direct causal role for cardenolides in *E. cheiranthoides* defense was recently demonstrated using CRISPR/Cas9 knockouts lacking cardenolides (Younkin et al. 2024). In a short-term field experiment, knockout plants showed a higher occurrence of snails and *P. rapae* eggs, whereas flea beetles and aphids were unaffected. Laboratory assays similarly revealed strong preference of *P. rapae* and some specialists for knockout plants (Younkin et al. 2024), while other specialists such as *P. xylostella* appear insensitive to cardenolides (Wang and Züst 2025). While these experiments establish the mechanistic basis of cardenolide resistance, our multi-year field study demonstrates how this resistance translates into reduced damage, altered growth trajectories, and context-dependent competitive outcomes under natural herbivore pressure.

Across all our analyses, the proportion of variance attributable to *E. cheiranthoides* genotype (intra-population variation) was moderate, with intraclass correlation coefficients (ICC) typically around 5%, but reaching up to 25% for individual responses (e.g., pollen beetle abundance). This degree of variation indicates substantial standing genetic diversity in growth- and defense-related traits among *E. cheiranthoides* genotypes. This variation could potentially enable future evolutionary tuning of defense allocation and cardenolide composition.

Concurrent production of glucosinolate and cardenolide defenses by *E. cheiranthoides* likely incurs substantial allocation costs. Both glucosinolates (in *Arabidopsis thaliana*) and cardenolide (in *Asclepias syriaca*) production can reduce growth under some conditions (Züst et al. 2011, 2015). Under low herbivory, such costs may reduce competitive ability, explaining inferior performance of *E. cheiranthoides* when herbivores were suppressed. In contrast, high resistance confers a clear advantage under high herbivory, as preferential attack on susceptible competitors indirectly favors the defended species. Such dynamics could shift species frequencies under sustained herbivory (Crawley 1989, Myers and Sarfraz 2017).

Despite weak competitive ability in the absence of specialized herbivores, the genus *Erysimum* has undergone a recent radiation that coincides with the gain of cardenolides (Züst et al. 2020), suggesting that an escape from herbivory aided diversification in enemy-free space. Even so, *Erysimum* species typically occupy marginal habitats and rarely dominate communities. Further evolutionary tuning may reduce defense for these plants in the future (e.g., Karasov et al. 2017), but increased *Erysimum* abundance could also intensify selection for cardenolide resistance in herbivores (Dobler et al. 2012, Petschenka et al. 2017), thus perpetuating the arms race between plants and their herbivores.

In conclusion, the gain of a novel defense appears to have allowed *Erysimum* plants to escape herbivore-imposed selection – if not herbivory entirely. However, maintaining this enemy-free space is costly, such that *Erysimum* thrives primarily where competitors are scarce or suppressed by herbivores. As these specific conditions are rare, *Erysimum* is currently largely confined to survive in marginal habitats, yet changing environmental conditions might well favor this unusually well-defended plant in the future.

## Acknowledgements

We thank Mark Charran, Broti Biswas, Teresa Vaello, Daniel Schlagenhauf, Alina Pfammatter and Paul Wieduwilt for field assistance, and Laura Dällenbach, Rayko Jonas and Markus Meierhofer for greenhouse support. We are grateful to Agroscope Reckenholz for the use of their field sites and infrastructure, and we especially thank Daniel Fuchs and Friedrich Käser for invaluable assistance in the field, and Robert Baur for administrative support. This project has received funding from the European Research Council (ERC) under the European Union’s Horizon 2020 research and innovation program (grant agreement No 950319). Additional support was provided by a Swiss National Science Foundation grant PCEFP3_194590 to TZ and a grant of the Georges and Antoine Claraz Foundation.

## Author Contributions

TZ conceived the project, KW and TZ designed the methodology, and all authors performed the experiments and analyzed the data. KW wrote the first draft of the manuscript, and all authors contributed critically to the drafts and gave final approval for publication.

## Supplementary Materials

### Supplementary Methods

#### 1. Analysis of herbivore community composition

We first analysed total herbivore abundance using separate negative binomial generalized linear mixed effects models (GLMMs, function *glmmTMB* in R package glmmTMB; Brooks et al., 2017) for each year. For each herbivore species, counts were summed per plant across six observation dates and modelled with fixed effects of plant species, herbivore species, and their interaction, with random intercepts included for plant genotype and plant identity. For the 2023 experiment, we excluded the competition treatment due to incomplete observations, but we included the insecticide treatment as an additional fixed effects with all interactions. Significance of fixed effects was assessed using Type III Wald Χ^2^ tests implemented in the function *Anova* (R package car). Post hoc pairwise comparisons were performed using function *emmeans* (R package emmeans) to test for plant-species differences under ambient conditions and insecticide effects within each plant species.

Next, we compared herbivore community composition for *R. nigrum* and *E. cheiranthoides* across years and among treatments (for the 2023 experiment) using multivariate analysis of field observation data. For comparisons between years, we excluded the competition and insecticide treatments of the 2023 experiment and summed observations from week 4 and 5 for each remaining plant (corresponding to peak herbivore abundance in both years). For comparison among treatments within in the 2023 experiment, we used data from week 4 only, as later weeks lacked observations for the competition treatment. For both reduced datasets, we calculated Bray-Curtis dissimilarity matrices from square-root-transformed herbivore counts and tested for differences in community composition using PERMANOVA (function *adonis2*, R package vegan; Oksanen et al. 2025). Homogeneity of multivariate dispersion was evaluated using function *permutest* applied to *betadisper* results, and non-metric multidimensional scaling (NMDS, function *metaMDS*) was used to visualize patterns in community composition (all in R package vegan).

#### 2. Analysis of temporal variation in herbivore abundances

Temporal variation in presence and abundance of the eight main herbivores was analyzed using generalized additive models (GAMs). We first fitted binomial GAMs to presence-absence data across all six weeks. We then excluded weeks with zero observations for each herbivore and fitted negative binomial GAMs to abundance data from the remaining weeks. For 2022, we constructed pairs of nested GAMs including plant species as a fixed effect but differing in their temporal smoothers: one model assuming a shared temporal pattern across plant species, and the other allowing plant-specific smoothers. Both models included random intercepts for plant genotype and plant identity nested within genotype. For binomial GAMs, the number of basis dimensions (parameter *k*) was fixed at 5, whereas for negative binomial GAMs, *k* was set to *W_pres_* – 1, where *W_pres_* denotes the number of observation weeks with herbivore presence. Nested models differing in the smoothing structure were fitted using maximum likelihood (ML) and compared via Akaike’s Information Criterion (AIC), preferring the simpler model unless ΔAIC was >2. The optimal model was then refitted using restricted maximum likelihood (REML), and significance of fixed effects was evaluated using Wald tests on model deviance.

For 2023, we extended this approach to include insecticide-treated plants. To determine the optimal fixed-effects structure for each herbivore, we compared nested models including either additive or interactive effects of plant species and insecticide treatments, while allowing temporal smoothers to vary across all four factor combinations. We then fixed the optimal fixed-effects structure (based on AIC) and compared nested models differing in the temporal smoother (global, by plant species, by insecticide, or by their combination). Each selected model was finally refitted using REML.

#### 3. Prediction of total leaf number from size measurements

Total leaf number was recorded only for a subset of plants in 2023. To obtain leaf number estimates for all plants, we used a two-stage predictive modelling approach. All models were trained separately for *R. nigrum* and *E. cheiranthoides* because the two species differ strongly in morphology and leaf numbers. For *R. nigrum*, we used candidate predictors of observation week, plant height, and plant width, including all pairwise interactions. For *E. cheiranthoides*, branch number and its pairwise interactions were additionally included as predictors because this species was frequently branched. Main effects and interactions were encoded using species-specific model-matrix representations, ensuring consistent predictor structure across all models within each species.

We used Poisson LASSO (least absolute shrinkage and selection operator) regression to identify a parsimonious set of predictors (function *cv.glmnet* in R package glmnet; Friedman et al. 2010). For each species, we fitted a cross-validated model using the subset of plants with observed total leaf number. To obtain a stable and conservative predictor selection, we performed 200 non-parametric bootstrap resamples for each species, refitting the LASSO model to each bootstrap sample. For every model refit, we recorded all predictors with non-zero coefficients at the regularization value *lambda.min*. Predictors retained in ≥ 70% of all bootstrap replicates were considered stable and used for the final model. This ‘stability selection’ step reduces sensitivity to sampling variation and guards against overfitting.

Because count data often show overdispersion relative to a Poisson distribution, we re-fitted the final predictive model for each species using a negative binomial generalized linear model (function *glm.nb* in R package MASS, Venables & Ripley 2002). Each negative binomial model included only the predictors identified through stability selection. These final models were then used to predict total leaf number for all plants lacking observations.

For subsequent analyses of herbivore damage, we always used the observed leaf number when available. For plants lacking observations, we used the predicted leaf number unless the predicted value was lower than the total number of damaged leaves, in which case the number of damaged leaves was used as the total leaf count. This ensures biological plausibility and prevents damage estimates from exceeding total leaf number.

#### 4. Plant height growth modelling

Plant height was recorded as a simple measure of plant size during damage assessments in 2022 (weeks 3, 6, and 9), and weekly in 2023. In 2022, height was measured from the root base to the tallest point of a plant, including leaves. However, this approach introduced variability due to differences in leaf size and leaf angle. To obtain a more standardized measure of structural growth, we recorded plant height in 2023 as stem height, measured from the root base to the apical meristem.

We modelled plant height growth using a four-parameter logistic function of the form

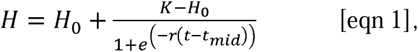

where plant height *H* at time *t* is a function of the initial plant height *H_0_* (i.e., height at transplanting), the asymptotic maximum height *K*, the growth rate *r*, and the time of the inflection point *t_mid_*(i.e., the time when height is halfway between *H_0_* and *K*). We fitted equation 1 to plant height data using the *nlme* function from the R package nlme, treating plant identity as random effect to account for repeated measures of individual plants.

As we only recorded height three times in the 2022 field experiment, our ability to estimate all four model parameters was limited. We therefore fixed parameters *t_mid_* and *r* at species-specific values, using an iterative process. For each species, we first set *r* = 0.1 and fitted models across a range of *t_mid_* values (20-60 days). We compared models using AIC and selected the value of *t_mid_*that minimized AIC. Holding *t_mid_* constant at this optimal value, we then repeated the procedure across a range of *r* values (0.05-0.5) to identify optimal species-specific values of *r* that minimized AIC. Using these optimal values as fixed parameters, we refitted the model to the full data to estimate the remaining two parameters, specifying fixed effects of plant species for both *H_0_* and *K*, and sequentially removing non-significant model terms.

For the 2023 field experiment, weekly recordings resulted in up to nine height measurements per plant. We removed plants with fewer than five observations (e.g., due to mortality), which allowed us to estimate all four parameters directly from the data. We also excluded plants from the competition treatment as we stopped recording their heights in the second half of the experiment. Fixed effects of plant species, insecticide treatment, and their interaction were included for parameters *K*, *t_mid_*, and *r*. For *H_0_*, we included only plant species as a fixed effect, as insecticide treatment was applied after transplanting and therefore could not influence initial height. Starting from a full model, we simplified the model by sequentially removing non-significant terms.

To compare growth dynamics between species and insecticide treatment over time, we calculated the absolute growth rate (AGR) from the derivative of equation 1:

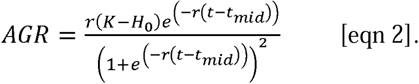

We calculated AGR for the midpoints of the first (weeks 3-6) and second (weeks 6-9) growth interval in both years. For each set of AGR estimates, we generated 95% population prediction intervals by drawing random values from the sampling distribution of each of the estimated parameters (Bolker, 2008).

#### 5. Leaf traits

Leaves from subsets of plants in the 2022 field experiment were harvested in weeks 4, 7, and 10 (one week after damage assessments) to assess temporal changes in plant nutritional quality. Each plant was only sampled once: plants from blocks 1-3 were sampled in week 4, blocks 4-6 in week 7, and blocks 7-10 in week 10. For each sampling, we removed 5-10 leaves from *R. nigrum* or 10-20 leaves from *E. cheiranthoides*, selecting and pooling leaves from the top, middle, and lower parts of each plant. Leaves from the same individual were wrapped together in aluminium foil and immediately placed on dry ice. Frozen samples were transported to the laboratory and stored at −80 °C before freeze-drying under vacuum for 48 h.

All *R. nigrum* samples were included in the analyses, whereas for *E. cheiranthoides* we analysed a random subset that also included genotypes excluded from the main analyses. For each selected plant, dried leaves were homogenized, and two aliquots of 10 mg were weighed into separate 2 mL screw-cap plastic tubes (Avantor, USA). One aliquot was used for determination of total protein content, and the other was used for determination of sugar and starch content.

##### 5.1 Total protein content

Total protein content of leaves was determined following Clissold et al. (2006). Briefly, 10 mg of ground leaf material was extracted twice using a total volume of 0.8 mL 0.1M NaOH. Samples were sonicated for 30 minutes, heated at 90 °C for 15 min, and centrifuged. Supernatants were pooled and neutralized with 6 M HCl. We precipitated proteins by adding 90 µL trichloroacetic acid solution (100% w/v, Sigma-Aldrich, Switzerland), followed by centrifugation and washing of the protein pellet with cold acetone. Dried pellets were resuspended in 1 mL of 0.1M NaOH, diluted 1:20 with water, and 50 µL of this solution was assayed photometrically using 200 µL QuickStart^TM^ Bradford Dye (BioRad, Switzerland). Absorbance was read at 595 nm on a microplate reader. Each sample was assayed in triplicate, and protein concentrations were calculated using an Immunoglobuline G (Sigma-Aldrich, Switzerland) standard curve (0-12 µg protein per assay).

##### 5.2 Soluble sugars and starch content

Soluble sugars were quantified following Machado et al. (2013). Ground plant material was extracted with 0.5 mL 80% ethanol, and re-extracted twice using 0.5 mL 80% ethanol. For each extraction, samples were vortexed, heated at 78 °C for 15 min, and centrifuged. Supernatants were pooled, and 25 µL of each extract were used for sequential determination of glucose, fructose, and sucrose using the enzymatic assay of Velterop & Vos (2001). Briefly, glucose was converted to glucose-6-phosphate by hexokinase (Roche, Switzerland) in the presence of ATP (Sigma-Aldrich, Switzerland), and further converted to 6-phosphogluconate by glucose-6-P-dehydrogenase (Roche, Switzerland) with NADP^+^ (Sigma-Aldrich, Switzerland). The resulting NADPH was quantified photometrically at 340 nm. After readings stabilized, phosphoglucoisomerase (Roche, Switzerland) was added to convert fructose to glucose, and absorbance was read again. Finally, we added invertase (Sigma-Aldrich, Switzerland) to convert sucrose into glucose and fructose, and absorbance was read a third time. All reactions were carried out in 50 mM HEPES buffer (pH 7, with 5 mM MgCl_2_). Each sample was run in triplicate, and sugars were quantified relative to a glucose standard curve (0-2 mM).

Starch content was determined from the pellet remaining after sugar extraction, following Smith & Zeeman (2006). We added 1 mL of water to each pellet and diluted 100 µL of this suspension with 500 µL water. Dilute suspensions were autoclaved for 1 h cooled to 37 °C. We then added 500 µL of an enzyme solution containing 50 mM sodium-acetate buffer (pH 5.5), amyloglucosidase (Roche, Switzerland) and α-amylase (Sigma-Aldrich, Switzerland). Samples were incubated overnight at 37 °C, and glucose equivalents were quantified in in 25 µL of supernatant using the same enzymatic assay as above.

##### 5.3 Trichome density and length

Trichomes were quantified on separate greenhouse-grown plants of *R. nigrum* and *E. cheiranthoides* (only using one genotype). Plants were germinated and grown as in the field experiments and maintained in the greenhouse for four weeks. We then measured trichome densities and lengths on both sides of single leaves from six plants per species. High-resolution images were acquired using an Emspira 3 digital microscope (Leica Microsystems, Switzerland, Figure S8a-d). Trichomes were counted within defined areas using ImageJ (Schneider et al. 2012). For three leaves per species and leaf side, we also measured the lengths of eight trichomes using ImageJ.

#### 6. Herbivore choice assays

To complement field observations with controlled tests of herbivore preference, compared feeding choices of four laboratory-reared herbivores between *R. nigrum* and *E. cheiranthoides* (only using one genotype). All used herbivores specialize on Brassicaceae and tolerate glucosinolates, but differ in their natural association with *E. cheiranthoides*. The turnip sawfly *Athalia rosae* (Hymenoptera: Tenthredinidae) has no known association with *E. cheiranthoides* but feeds on other Brassicaceae

plants as larvae and sequesters glucosinolates from its host plants (Müller & Wittstock, 2005). The cabbage white butterfly *Pieris rapae* (Lepidoptera: Pieridae) actively avoids *E. cheiranthoides* (Sachdev-Gupta et al., 1993; Younkin et al., 2024) but its caterpillars feed on other Brassicaceae plants by detoxifying activated glucosinolates enzymatically (Wittstock et al., 2004). The diamondback moth *Plutella xylostella* (Lepidoptera: Plutellidae) is occasionally found feeding on *E. cheiranthoides* (Mertens et al., 2021; observations in current study) and its caterpillars enzymatically degrade glucosinolates prior to their activation (Ratzka et al., 2002), while the cardenolide coping mechanism remains unknown. Finally, the red cabbage bug *Eurydema ornata* (Hemiptera: Pentatomidae) is frequently found feeding on *E. cheiranthoides* and other *Erysimum* species both as adults and nymphs (Goméz, 2005; observations in current study). Individuals sequester glucosinolates from their host plant (Aliabadi et al., 2002), but again the cardenolide coping mechanism for this species remains unknown.

Individuals of *A. rosae* were kindly provided by Prof. Caroline Müller, University of Bielefeld, and had been maintained as a laboratory rearing colony on *Sinapis arvensis*. Newly eclosed, unmated females were left to oviposit on *Brassica rapa* plants (Wisconsin Fast Plants^TM^, Caroline Biological Supply Co., Burlington VA, USA), and the developing male larvae were used for choice assays 5 days after hatching. Individuals of *P. rapae* were kindly provided by the group of Prof. Florian Schiestl, University of Zürich. Adults had originally been collected from multiple locations in Switzerland and were maintained in a panmictic laboratory rearing colony using *Brassica rapa* plants as oviposition substrate and leaves of savoy cabbage (*Brassica oleracea* var. *sabauda*) as caterpillar food. We placed fresh *B. rapa* plants in an adult oviposition cage for 24 hours, after which larvae were left to develop on the oviposition plant for 10 days before use in choice assays (early L2 stage). For *P. xylostella* we used a strain from a long-term rearing on a wheat germ-based artificial diet (Frontier Agricultural Sciences, Delaware, USA), originally provided by Syngenta Crop Protection AG, Stein, Switzerland. For choice assays, newly laid eggs on sheets of parafilm were placed in cups of fresh diet and left to develop for 11 days (late L2 stage). We originally collected adults of *E. ornata* from a population in Valais, Switzerland (46.312970° N, 7.675072° E, predominantly collected on *Isatis tinctoria*), and maintained a continuous laboratory rearing population on a diet of savoy cabbage leaves and young *B. rapa* plants, with yearly introductions of new individuals from the same wild population to maintain genetic diversity. For choice assays, we used mature adults (>3 days after final molt) out of the rearing population.

Choice assays were set up in petri dishes (5 cm diameter) containing a thin layer of 3% agar, with 12 replicates per herbivore species. Each petri dish contained paired leaf discs (7.6 mm diameter), one from *R. nigrum* and one from *E. cheiranthoides* (4-week old greenhouse plants). A single larva (for *A. rosae*, *P. rapae*, *P. xylostella*) or adult (for *E. ornata*) was released at dish centre. Larvae (leaf chewers) were allowed to feed for 6 h, whereas *E. ornata* (cell-content feeder) fed for 24 h. Remaining leaf discs were scanned using a CanoScan LiDE 220 scanner, and remaining leaf area was quantified in ImageJ. Consumed area was calculated by subtracting remaining leaf areas from the area of an intact leaf disc (45.36 mm^2^).

Consumed leaf areas were analysed with a linear mixed-effects model including plant species, herbivore species, and their interaction as fixed effects, and Petri dish ID as a random effect to account for paired discs. Species-specific preferences were then tested using pairwise contrasts (function *emmeans*).

### Supplementary Results

#### Herbivore community composition

We compared herbivore community composition between years by focused on individual plants under ambient herbivore conditions and summed observations from weeks 4 and 5, representing peak abundance in both years. Using PERMANOVA, we found a significant interaction between plant species and year (F_1,408_ = 10.32, p < 0.001), although group dispersion also differed substantially for both factors (permutation tests on *betadisper*, *p* < 0.001).

To compare composition among the factorial combination of competition and insecticide treatments in 2023, we analysed observations from week 4, corresponding to high abundance and complete sampling across all treatments. We detected a marginally significant three-way interaction between plant species, insecticide treatment, and competition (F_1,283_ = 2.49, p = 0.070), thus we analysed plants separately. For *R. nigrum*, community composition differed by insecticide (F_1,85_ = 22.45, p < 0.001) but not by competition (F_1,85_ = 1.28, p = 0.274), and group dispersion was homogeneous across treatment (permutation tests, insecticide: p = 0.680, competition: p = 0.119; Supplementary Figure S2a). In contrast, community composition on *E. cheiranthoides* differed significantly by both insecticide (F_1,200_ = 79.04, p < 0.001) and competition (F_1,200_ = 21.19, p < 0.001), but group dispersion also differed for the competition treatment (permutation tests, insecticide: p = 0.420, competition: p < 0.001; Supplementary Figure S2b).

#### Species differences in plant traits

In the 2022 field experiment, we quantified nutritional traits that may influence herbivore preference. Across three sampling points, *R. nigrum* leaves contained significantly higher concentrations of soluble glucose and fructose (Fig. S7a-b), whereas *E. cheiranthoides* contained higher levels of sucrose (Fig. S7c). Digestible starch content did not differ between species for most of the experiment, except at the final sampling, when starch had declined in both species but more strongly in *E. cheiranthoides* (Fig. S7d). Total hydrolysable protein content differed only at later time points, with *R. nigrum* containing up to 60% more protein per unit dry mass.

Trichomes differed markedly between the two plant species, with *R. nigrum* producing long, unbranched trichomes (Figure S8a-b), whereas *E. cheiranthoides* had smaller, medifixed stellate trichomes (Figure S8c-d). Trichome density on *R. nigrum* was lowest on the upper (adaxial) surface (0.04 ± 0.01 trichomes mm^-2^; mean ± 1 SE) and slightly higher on the lower (abaxial) surface (0.21 ± 0.07 trichomes mm^-2^), where trichomes were predominantly located along veins. In contrast, *E. cheiranthoides* had very high trichome densities on both surfaces (adaxial: 8.84 ± 0.61 trichomes mm^-2^; abaxial: 7.50 ± 0.52 trichomes mm^-2^). Although trichome density and morphology may vary with plant age and environment, these measurements nonetheless provide a useful reference for the relative differences between *R. nigrum* and *E. cheiranthoides*.

---

**Table S1.** Mean (± 1 SE) numbers of herbivores per plant observed sitting or feeding on *R. nigrum* and *E. cheiranthoides* in the 2022 field experiment, summed across six observation dates. Values are back-transformed model predictions from a negative binomial generalized linear mixed-effects model of herbivore counts as a function of plant species and herbivore species. Numbers in parentheses in the table header indicate the number of replicate plants per species. Pairwise linear contrasts were used to test for differences between plants for each herbivore species (*Rha vs. Ery*). Bold values highlight significant contrasts. Flea beetles, large flea beetles, weevils, and leaf hoppers refer to taxonomically complex groups that were not identified to species level in the field.

| Herbivore | Scientific name | Order | Stage | *R. nigrum* (n = 28) | *E. cheiranthoides* (n = 62) | Pairwise contrast<br>*Rha* vs. *Ery* |
| --- | --- | --- | --- | --- | --- | --- |
| flea beetle | | Coleoptera | adult | 24.10 ± 4.50 | 1.61 ± 0.26 | **z-ratio = 11.10, p < 0.001** |
| pollen beetle | *Brassicogethes aeneus* | Coleoptera | adult | 88.20 ± 16.00 | 4.95 ± 0.68 | **z-ratio = 12.70, p < 0.001** |
| weevil | | Coleoptera | adult | 3.94 ± 0.69 | 0.81 ± 0.13 | **z-ratio = 6.22, p < 0.001** |
| leaf miner | *Phytomyza* sp. | Diptera | juvenile | 0.34 ± 0.12 | 0.04 ± 0.03 | **z-ratio = 2.96, p = 0.003** |
| aphid | | Hemiptera | adult/juvenile | 6.92 ± 1.39 | 0.66 ± 0.13 | **z-ratio = 8.35, p < 0.001** |
| cabbage whitefly | *Aleyrodes* sp. | Hemiptera | adult | 7.74 ± 1.52 | 0.29 ± 0.07 | **z-ratio = 10.2, p < 0.001** |
| leaf hopper | | Hemiptera | adult | 0.06 ± 0.05 | 0.06 ± 0.03 | z-ratio = 0.07, p = 0.943 |
| crucifer shield bug | *Eurydema* sp. | Heteroptera | adult/juvenile | 0.64 ± 0.19 | 0.70 ± 0.14 | z-ratio = -0.29, p = 0.775 |
| plant bug | *Lygus* sp. | Heteroptera | adult/juvenile | 5.12 ± 1.02 | 0.83 ± 0.15 | **z-ratio = 6.68, p < 0.001** |
| potato capsid | *Closterotomus norwegicus* | Heteroptera | adult | 0.16 ± 0.08 | 0.07 ± 0.03 | z-ratio = 1.13, p = 0.260 |
| diamondback moth | *Plutella xylostella* | Lepidoptera | adult/juvenile | 1.70 ± 0.40 | 0.59 ± 0.12 | **z-ratio = 3.46, p < 0.001** |

**Table S2.**
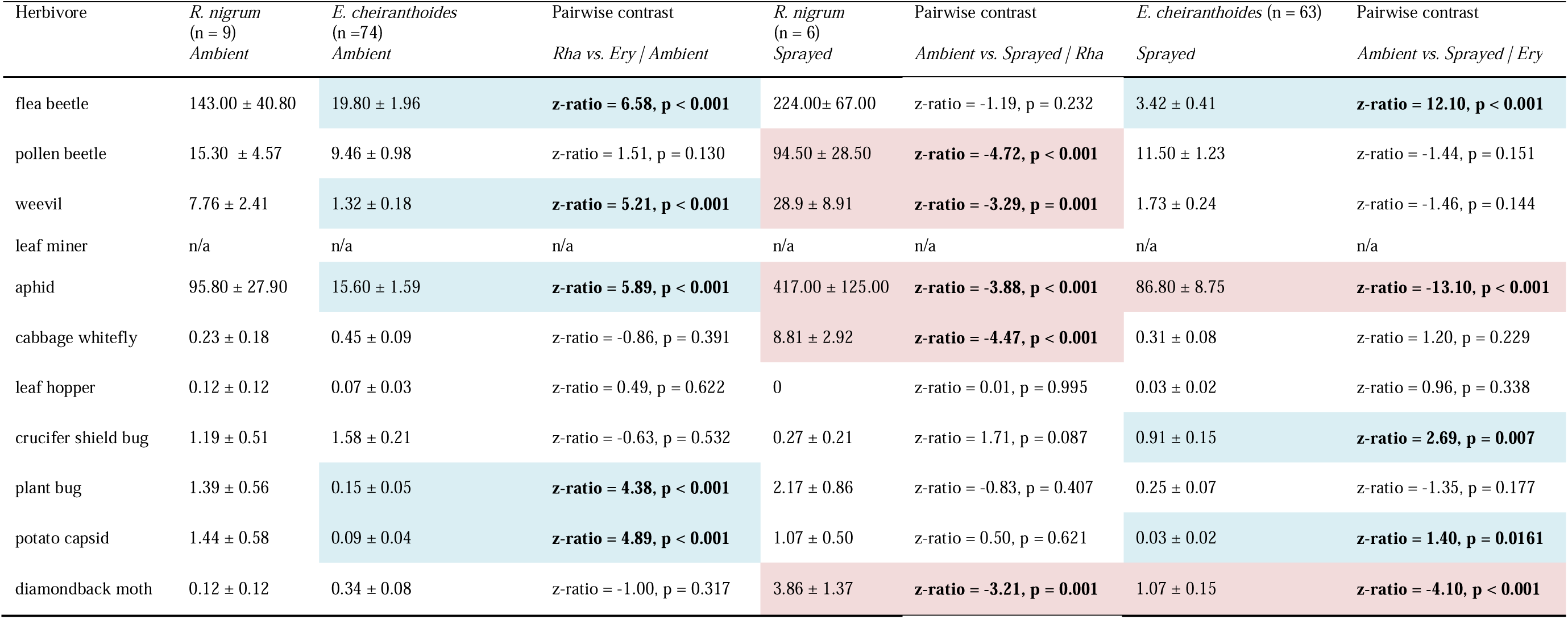
Mean (± 1 SE) numbers of herbivores per plant observed sitting or feeding on individual *R. nigrum* and *E. cheiranthoides* in the 2023 field experiment (excluding plants of the competition treatment), summed across six observation dates. Values are back-transformed model predictions from a negative binomial generalized linear mixed-effects model of herbivore counts as a function of plant species, herbivore species, and insecticide treatment. Numbers in parentheses in the table header indicate the number of replicate plants per species and treatment. Pairwise linear contrasts were used to test for differences between plant species under ambient conditions (*Rha vs. Ery | Ambient)*, and for the effects of insecticide treatment within *R. nigrum* (*Ambient vs. Sprayed | Rha*) and *E. cheiranthoides* (*Ambient vs. Sprayed | Ery*). Bold values highlight significant contrasts, and colours additionally highlight significant decreases (blue) or increases (red) in mean numbers for each contrast. Leaf miners were never observed on *R. nigrum* in 2023 and could not be analysed in this model.

| Herbivore | *R. nigrum*<br>(n = 9)<br>*Ambient* | *E. cheiranthoides*<br>(n = 74)<br>*Ambient* | Pairwise contrast<br>*Rha* vs. *Ery* / *Ambient* | *R. nigrum*<br>(n = 6)<br>*Sprayed* | Pairwise contrast<br>*Ambient* vs. *Sprayed* / *Rha* | *E. cheiranthoides* (n = 63)<br>*Sprayed* | Pairwise contrast<br>*Ambient* vs. *Sprayed* / *Ery* |
| --- | --- | --- | --- | --- | --- | --- | --- |
| flea beetle | 143.00 ± 40.80 | 19.80 ± 1.96 | **z-ratio = 6.58, p < 0.001** | 224.00 ± 67.00 | z-ratio = -1.19, p = 0.232 | 3.42 ± 0.41 | **z-ratio = 12.10, p < 0.001** |
| pollen beetle | 15.30 ± 4.57 | 9.46 ± 0.98 | z-ratio = 1.51, p = 0.130 | 94.50 ± 28.50 | **z-ratio = -4.72, p < 0.001** | 11.50 ± 1.23 | z-ratio = -1.44, p = 0.151 |
| weevil | 7.76 ± 2.41 | 1.32 ± 0.18 | **z-ratio = 5.21, p < 0.001** | 28.9 ± 8.91 | **z-ratio = -3.29, p = 0.001** | 1.73 ± 0.24 | z-ratio = -1.46, p = 0.144 |
| leaf miner | n/a | n/a | n/a | n/a | n/a | n/a | n/a |
| aphid | 95.80 ± 27.90 | 15.60 ± 1.59 | **z-ratio = 5.89, p < 0.001** | 417.00 ± 125.00 | **z-ratio = -3.88, p < 0.001** | 86.80 ± 8.75 | **z-ratio = -13.10, p < 0.001** |
| cabbage whitefly | 0.23 ± 0.18 | 0.45 ± 0.09 | z-ratio = -0.86, p = 0.391 | 8.81 ± 2.92 | **z-ratio = -4.47, p < 0.001** | 0.31 ± 0.08 | z-ratio = 1.20, p = 0.229 |
| leaf hopper | 0.12 ± 0.12 | 0.07 ± 0.03 | z-ratio = 0.49, p = 0.622 | 0 | z-ratio = 0.01, p = 0.995 | 0.03 ± 0.02 | z-ratio = 0.96, p = 0.338 |
| crucifer shield bug | 1.19 ± 0.51 | 1.58 ± 0.21 | z-ratio = -0.63, p = 0.532 | 0.27 ± 0.21 | z-ratio = 1.71, p = 0.087 | 0.91 ± 0.15 | **z-ratio = 2.69, p = 0.007** |
| plant bug | 1.39 ± 0.56 | 0.15 ± 0.05 | **z-ratio = 4.38, p < 0.001** | 2.17 ± 0.86 | z-ratio = -0.83, p = 0.407 | 0.25 ± 0.07 | z-ratio = -1.35, p = 0.177 |
| potato capsid | 1.44 ± 0.58 | 0.09 ± 0.04 | **z-ratio = 4.89, p < 0.001** | 1.07 ± 0.50 | z-ratio = 0.50, p = 0.621 | 0.03 ± 0.02 | **z-ratio = 1.40, p = 0.0161** |
| diamondback moth | 0.12 ± 0.12 | 0.34 ± 0.08 | z-ratio = -1.00, p = 0.317 | 3.86 ± 1.37 | **z-ratio = -3.21, p = 0.001** | 1.07 ± 0.15 | **z-ratio = -4.10, p < 0.001** |

**Table S3.** Generalized additive model (GAM) results for temporal dynamics in herbivore presence in the 2022 and 2023 field experiments. a) For 2022, separate binomial GAMs were fitted to presence-absence data of each herbivore with a fixed effect of plant species. Nested models with either a global temporal smoother (‘day’), plant-specific smoothers (‘day by plant’) were compared based on AIC, retaining the simpler model unless ΔAIC >2. The number smoother nodes (*k*) were fixed to 5 for each model, and significance of fixed effect was assessed using Type III Wald chisquare tests on explained deviance. b) For 2023, equivalent GAMs were fitted with fixed effects of plant species and insecticide treatment (plants of the competition treatment were excluded). Models with additive *vs.* interactive fixed effects were compared based on AIC, followed by comparison of alternative smoother structures (‘day’, ‘day by plant’, ‘day by insecticide’, ‘day by plant × insecticide’). No model could be fitted for crucifer shield bugs on insecticide-treated plants due to insufficient observations.

| Herbivore | Plant |  | Smoother |
| --- | --- | --- | --- |
| aphids | **df = 1, $X^2 = 36.77$** | **p < 0.001** | day |
| cabbage whitefly | **df = 1, $X^2 = 39.25$** | **p < 0.001** | day |
| crucifer shield bug | df = 1, $X^2 = 1.10$ | p = 0.295 | day |
| diamondback moth | **df = 1, $X^2 = 19.66$** | **p < 0.001** | day |
| flea beetle | **df = 1, $X^2 = 5.78$** | **p = 0.016** | day by plant |
| plant bug | df = 1, $X^2 = 2.16$ | p = 0.142 | day by plant |
| pollen beetle | **df = 1, $X^2 = 37.86$** | **p < 0.001** | day by plant |
| weevil | **df = 1, $X^2 = 21.37$** | **p < 0.001** | day by plant |

b) 2023 field experiment
| Herbivore | Plant | | Insecticide | | Plant × Insecticide | | Smoother |
| --- | --- | --- | --- | --- | --- | --- | --- |
| aphids | df = 1, $X^2 = 0.05$ | p = 0.829 | **df = 1, $X^2 = 18.32$** | **p < 0.001** | df = 1, $X^2 = 2.20$ | p = 0.138 | day by insecticide |
| cabbage whitefly | df = 1, $X^2 = 0.95$ | p = 0.331 | **df = 1, $X^2 = 5.90$** | **p = 0.015** | **df = 1, $X^2 = 6.25$** | **p = 0.013** | day |
| crucifer shield bug | df = 1, $X^2 = 1.38$ | p = 0.241 | n/a | n/a | n/a | n/a | day by plant |
| diamondback moth | df = 1, $X^2 = 0.96$ | p = 0.327 | **df = 1, $X^2 = 8.37$** | **p = 0.004** | **df = 1, $X^2 = 4.63$** | **p = 0.031** | day |
| flea beetle | **df = 1, $X^2 = 27.51$** | **p < 0.001** | **df = 1, $X^2 = 46.22$** | **p < 0.001** | n.s. | n.s. | day by insecticide |
| plant bug | df = 1, $X^2 = 0.04$ | p = 0.834 | df = 1, $X^2 = 3.17$ | p = 0.075 | n.s. | n.s. | day by plant |
| pollen beetle | **df = 1, $X^2 = 6.34$** | **p = 0.012** | **df = 1, $X^2 = 5.72$** | **p = 0.017** | **df = 1, $X^2 = 5.61$** | **p = 0.018** | day by plant |
| weevil | **df = 1, $X^2 = 11.24$** | **p < 0.001** | df = 1, $X^2 = 3.04$ | p = 0.081 | df = 1, $X^2 = 1.19$ | p = 0.276 | day |

**Table S4.** Generalized additive model (GAM) results for temporal dynamics in herbivore abundance in the 2022 and 2023 field experiments. a) For 2022, separate negative binomial GAMs were fitted to count data of each herbivore after weeks with zero observations were excluded. Models included fixed effect of plant species, and nested models with a global temporal smoother (‘day’) or a plant-specific smoothers (‘day by plant’) were compared based on AIC, retaining the simpler model unless ΔAIC >2. The number smoother nodes (*k*) was adjusted for each model according to the number of weeks in which herbivores were observed. Significance of fixed effect was assessed using Type III Wald chisquare tests on explained deviance, and intraclass correlation coefficients (ICC) were calculated to evaluate plant genotype effects. No model could be fitted for cabbage whiteflies due to insufficient observations. b) For 2023, equivalent GAMs were fitted with fixed effects of plant species and insecticide treatment (plants of the competition treatment were excluded). Models with additive *vs.* interactive fixed effects were compared based on AIC, followed by comparison of alternative smoother structures. No model could be fitted for crucifer shield bugs on insecticide-treated plants due to insufficient observations.

| Herbivore | Plant |  | Smoother | k | ICC Genotype |
| --- | --- | --- | --- | --- | --- |
| aphids | **df = 1, $X^2 = 68.47$** | **p < 0.001** | day by plant | 5 | 0.0000 |
| crucifer shield bug | df = 1, $X^2 = 0.11$ | p = 0.740 | day | 3 | 0.0000 |
| diamondback moth | **df = 1, $X^2 = 26.36$** | **p < 0.001** | day | 3 | 0.0000 |
| flea beetle | **df = 1, $X^2 = 35.96$** | **p < 0.001** | day by plant | 5 | 0.0740 |
| plant bug | **df = 1, $X^2 = 26.89$** | **p < 0.001** | day by plant | 3 | 0.0464 |
| pollen beetle | **df = 1, $X^2 = 171.14$** | **p < 0.001** | day by plant | 3 | 0.0208 |
| weevil | **df = 1, $X^2 = 19.06$** | **p < 0.001** | day by plant | 3 | 0.0901 |

b) 2023 field experiment
| Herbivore | Plant |  | Insecticide |  | Plant × Insecticide |  | Smoother | k | ICC Genotype |
| --- | --- | --- | --- | --- | --- | --- | --- | --- | --- |
| aphids | **df = 1, $X^2 = 13.62$** | **p < 0.001** | **df = 1, $X^2 = 78.31$** | **p < 0.001** | n.s. | n.s. | day by plant × insecticide | 5 | 0.0636 |
| cabbage whitefly | df = 1, $X^2 = 0.44$ | p = 0.509 | **df = 1, $X^2 = 12.58$** | **p < 0.001** | **df = 1, $X^2 = 15.10$** | **p < 0.001** | day by insecticide | 3 | 0.0277 |
| crucifer shield bug | df = 1, $X^2 = 0.70$ | p = 0.403 | n/a | n/a | n/a | n/a | day by plant | 3 | 0.1378 |
| diamondback moth | df = 1, $X^2 = 1.01$ | p = 0.314 | **df = 1, $X^2 = 9.35$** | **p = 0.002** | **df = 1, $X^2 = 4.57$** | **p = 0.032** | day | 4 | 0.0290 |
| flea beetle | **df = 1, $X^2 = 126.89$** | **p < 0.001** | df = 1, $X^2 = 0.09$ | p = 0.769 | **df = 1, $X^2 = 37.74$** | **p < 0.001** | day | 5 | 0.0000 |
| plant bug | **df = 1, $X^2 = 39.57$** | **p < 0.001** | df = 1, $X^2 = 2.24$ | p = 0.134 | n.s. | n.s. | day by plant | 3 | 0.0000 |
| pollen beetle | df = 1, $X^2 = 0.00$ | p = 0.987 | **df = 1, $X^2 = 18.46$** | **p < 0.001** | **df = 1, $X^2 = 5.61$** | **p < 0.001** | day by plant × insecticide | 3 | 0.2579 |
| weevil | **df = 1, $X^2 = 23.08$** | **p < 0.001** | **df = 1, $X^2 = 6.78$** | **p = 0.009** | n.s. | n.s. | day | 3 | 0.1144 |

**Table S5.** a) Generalized-linear mixed effects model (GLMM) for the proportion of damaged leaves in the 2022 field experiment. Week, plant species, and their interaction were included as fixed effects. b) GLMM for the proportion of highly damaged leaves in the 2023 field experiment. Week, plant species, insecticide treatment, competition treatment, and all possible interactions were initially included as fixed effects. Because leaf damage was not recorded in week 9, interactions between competition treatment and week could not be fitted. A non-significant interaction of plant species × competition was removed from the full model, and statistics for the remaining terms are reported from the reduced model. An additive fixed effect of observer was included in all models.

|  | Df | Chisq | p-value |
| --- | --- | --- | --- |
| Week | 2 | 115.10 | < **0.001** |
| Plant | 1 | 168.91 | < **0.001** |
| Week:Plant | 2 | 54.17 | < **0.001** |
| Observer | 3 | 92.11 | < **0.001** |

Model: glmer(cbind(damaged,undamaged) ~ Week \* Plant + Observer + (1|Plant ID) + (1| Genotype) b) Analysis of Deviance Table (Type III Wald Chisquare tests)
|  | Df | Chisq | p-value |
| --- | --- | --- | --- |
| Week | 2 | 65.64 | < **0.001** |
| Plant | 1 | 126.52 | < **0.001** |
| Insecticide | 1 | 0.55 | 0.457 |
| Competition | 1 | 8.09 | **0.004** |
| Week:Plant | 2 | 19.91 | < **0.001** |
| Week:Insecticide | 2 | 39.06 | < **0.001** |
| Week:Plant:Insecticide | 2 | 94.42 | < **0.001** |
| Plant:Insecticide | 1 | 44.67 | < **0.001** |
| Plant:Competition | 1 | 1.88 | 0.171 |
| Insecticide:Competition | 1 | 12.18 | < **0.004** |
| Observer | 1 | 60.38 | < **0.001** |

Model: glmer(cbind(highly damaged, low+undamaged) ~ Week \* Plant \* Insecticide + Competition + Plant:Competition + Insecticide:Competition + Observer + (1|Plant ID) + (1|Neighbour group) + (1|Genotype)

**Table S6.** Parameters of four-parameter logistic models of plant height growth. a) 2022 field experiment: Because plant height was only recorded three times, only two parameters could be estimated from the data. We therefore compared a series of models across biologically relevant ranges of *t_mid_* and *r* for each species using AIC. After identifying optimal species-specific values for *t_mid_* and *r* set, we refitted the full model to estimate *H_0_* and *K* using an *nlme* model. Reported parameters are either model estimates with 95% confidence intervals or fixed optimized values. The negative estimate of *H_0_*for *E. cheiranthoides* (implying negative height at transplanting) likely reflects limited data on early development. *H_0_*, differed significantly by plant species, whereas there was no statistical support for plant-specific variation in *K*. b) 2023 field experiment: Weekly stem height measurements allowed the estimation of all four parameters using an *nlme* model. Reported parameters are model estimates with 95% confidence intervals. All parameters differed significantly between plant species, and *K* additionally varied with the plant species × insecticide interaction.

| a) 2022 field experiment |  |  |  |  |
| --- | --- | --- | --- | --- |
|  | *R. nigrum* | *E. cheiranthoides* |  |  |
| $H_0$ (estimated) | 19.56 [18.26 – 20.85] | -3.82 [-4.79 – -2.84] | | |
| $K$ (estimated) | 84.6 ± 1.91 | | | |
| $t_{mid}$ (fixed) | 56 | 36 | | |
| $r$ (fixed) | 0.10 | 0.09 | | |
| b) 2023 field experiment |  |  |  |  |
|  | *R. nigrum* |  | *E. cheiranthoides* |  |
|  | Ambient | Insecticide | Ambient | Insecticide |
| $H_0$ (estimated) | 1.65 [1.29 – 2.01] | | -0.23 [-0.39 – -0.08] | |
| $K$ (estimated) | 26.8 [18.3 – 35.3] | 55.8 [46.9 – 64.8] | 69.1 [66.0 – 72.1] | 75.5 [72.4 – 78.6] |
| $t_{mid}$ (estimated) | 38.0 [35.8 – 40.2] | | 39.4 [38.6 – 40.1] | |
| $r$ (estimated) | 0.283 [0.230 – 0.369] | | 0.115 [0.111 – 0.118] | |

**Table S7.** a) Linear mixed effects model (LMM) for ln-transformed biomass of plants in the 2022 field experiment, harvested in weeks 8, 11, and 14. Week, plant species, and their interaction were included as fixed effects. b) LMMs for ln-transformed biomass of plants in the 2023 field experiment, all harvested in week 10. Plant species, insecticide treatment, competition treatment, and all interactions were included as fixed effects.

|  | Df | Chisq | p-value |
| --- | --- | --- | --- |
| Week | 2 | 27.78 | **< 0.001** |
| Plant | 1 | 9.11 | **0.003** |
| Week:Plant | 2 | 6.49 | **0.039** |

Model: lmer(ln(biomass) ~ Week \* Plant + (1| Genotype) b) Analysis of Deviance Table (Type III Wald Chisquare tests)
|  | Df | Chisq | p-value |
| --- | --- | --- | --- |
| Plant | 1 | 87.11 | **< 0.001** |
| Insecticide | 1 | 33.45 | **< 0.001** |
| Competition | 1 | 13.40 | **< 0.001** |
| Plant:Insecticide | 1 | 25.46 | **< 0.001** |
| Plant:Competition | 1 | 5.83 | **0.016** |
| Insecticide:Competition | 1 | 4.64 | **0.031** |
| Plant:Insecticide:Competition | 1 | 14.52 | **< 0.001** |

Model: lmer(ln(biomass) ~ Plant \* Insecticide \* Competition + (1|Genotype)

**Figure S1.**
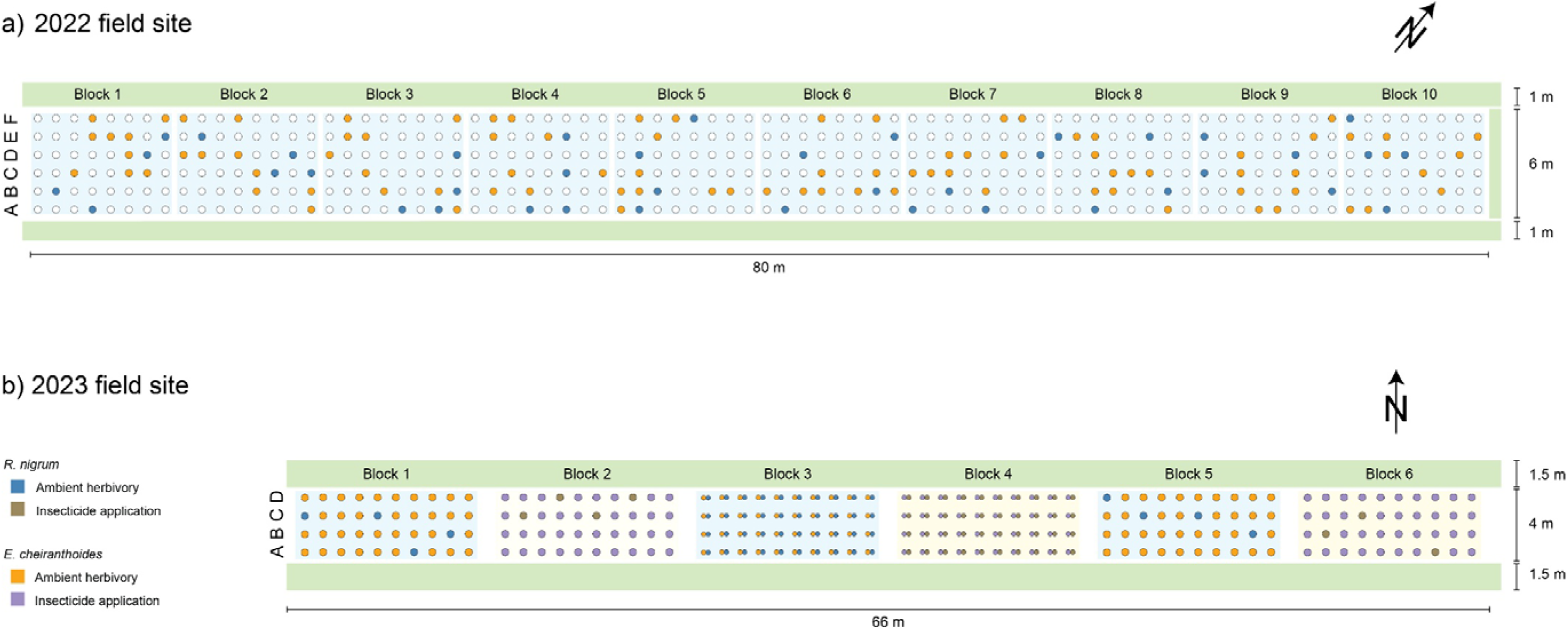
a) Schematic of the 2022 field site. Shown are the ten replicate blocks (light blue squares), with focal *R. nigrum* and *E. cheiranthoides* plants highlighted in colours. Empty circles are additional *E. cheiranthoides* plants belonging to genotypes not included for the analyses of this study. b) Schematic of the 2023 field site. Shown are the three blocks under ambient herbivory (light blue squares), and the three blocks receiving weekly applications of insecticide (light yellow squares). Paired circles in blocks 3 and 4 represent pairs of *R. nigrum* and *E. cheiranthoides* plants growing closely together in the same buried pot to impose plant-plant competition. Buffer strips sown with a mixture of *Raphanus sativus* and *Avena strigosa* are shown as light green rectangles for both years. Both sites were located adjacent to oilseed rape crop fields (not shown).

**Figure S2.**
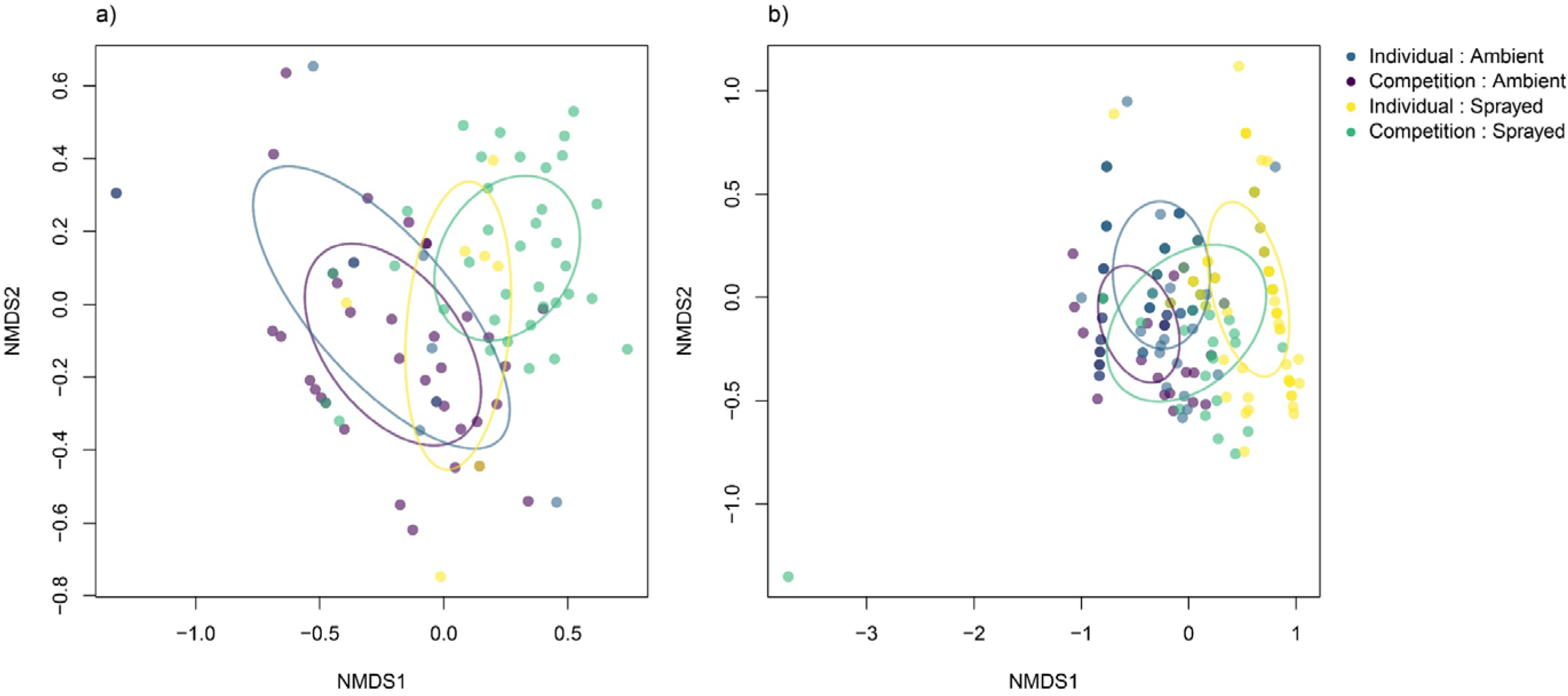
NMDS plots of the observed herbivore communities on a) *R. nigrum* or b) *E. cheiranthoides* in week 4 of the 2023 field experiment. Points are individual plants, with colours corresponding to the four combinations of the competition and insecticide treatment. Ellipses represent the within-treatment dispersion and show one standard deviation around the group centroid in NMDS space.

**Figure S3.**
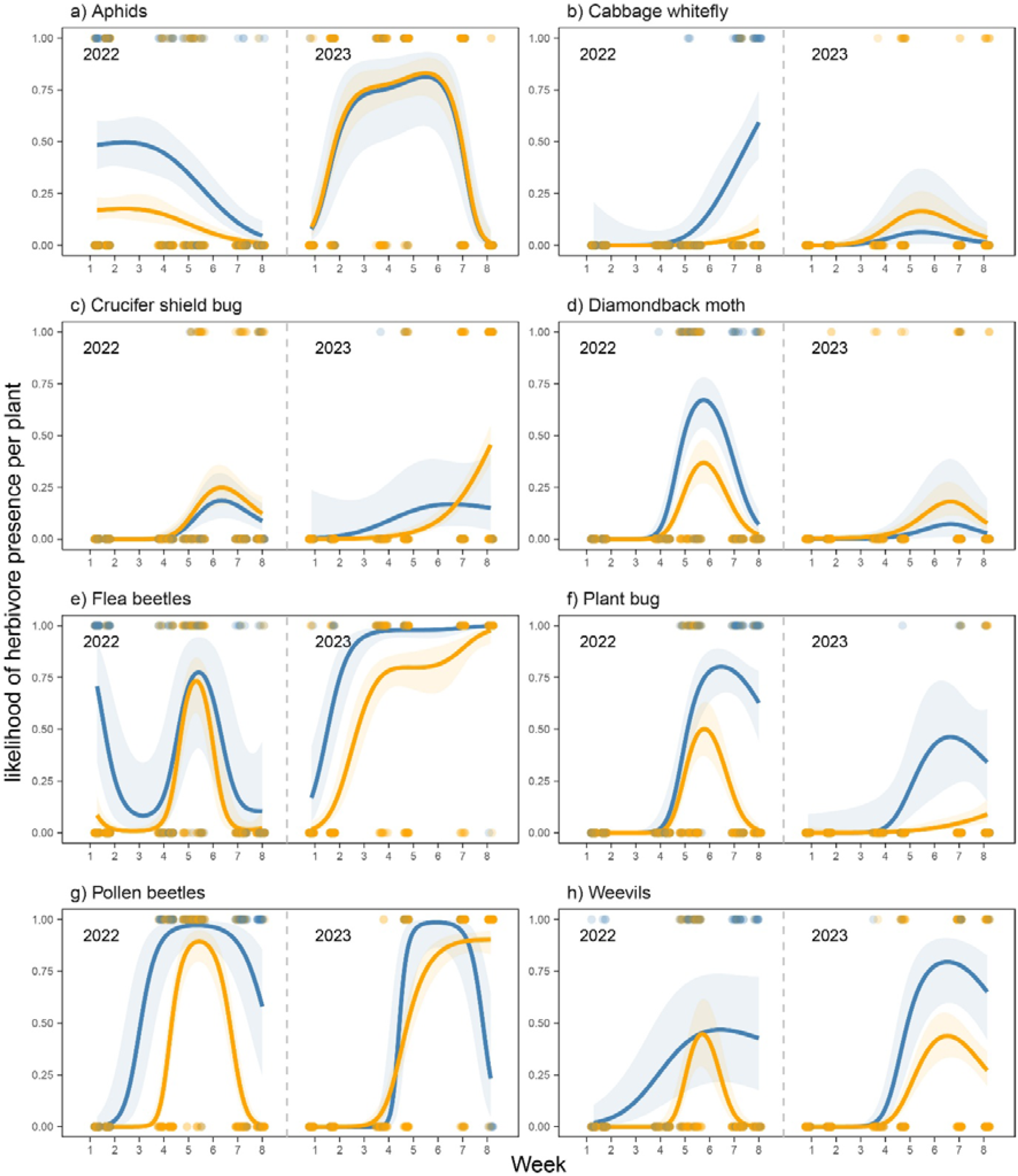
Likelihood of herbivore presence on *R. nigrum* (blue) and *E. cheiranthoides* (orange) in the 2022 and 2023 field experiment. Points are observations on individual plants, while lines and shaded are the mean model predictions and 95% confidence intervals from binomial generalized additive models (GAMs). All models used ‘days in the field’ as temporal variable, but the x-axes show ‘week’ for easier comparison with other results. See Supplementary Table S3 for test statistics of each model.

**Figure S4.**
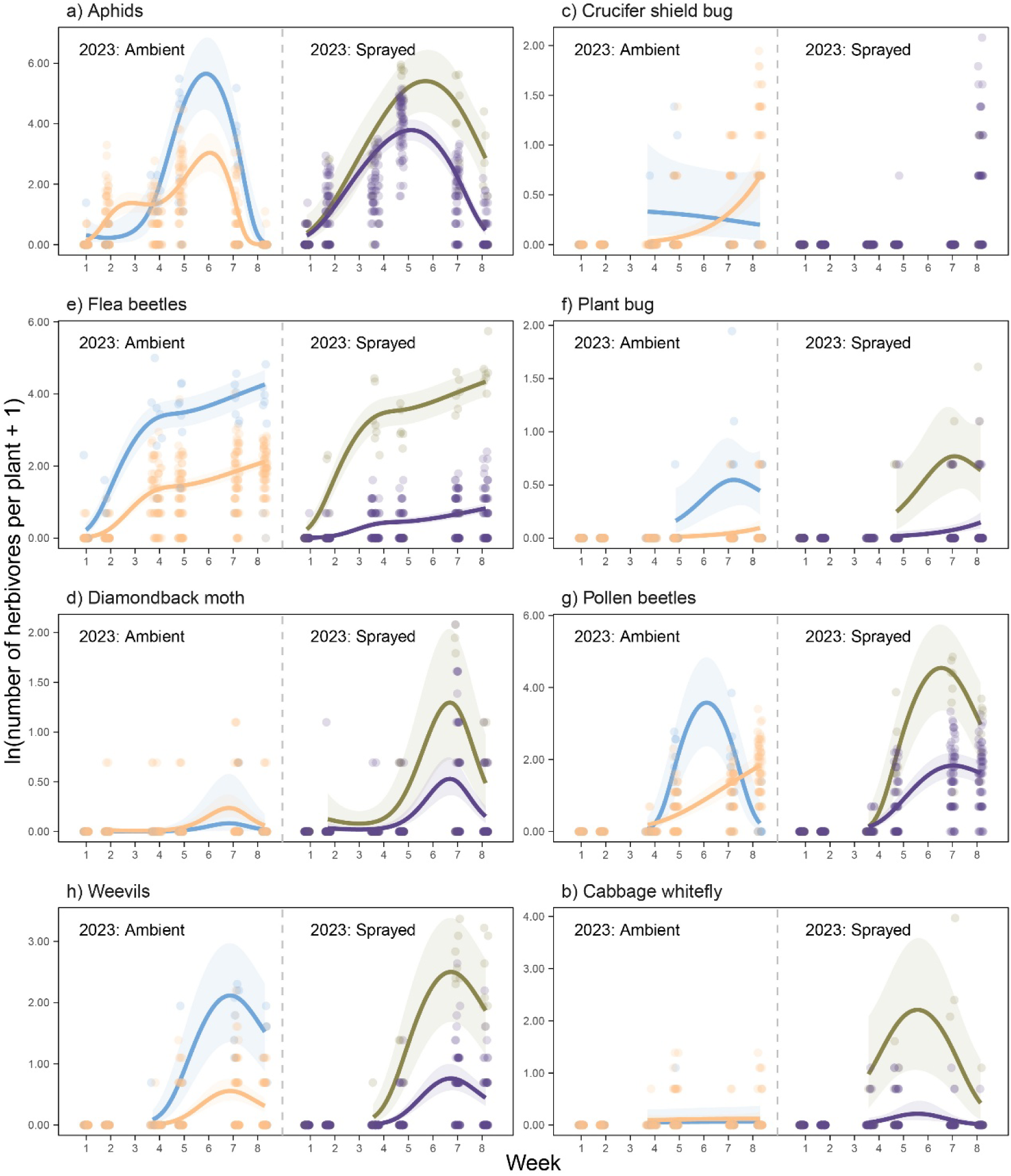
Herbivore abundance for the eight main herbivores under ambient or insecticide-treated conditions on individual *R. nigrum* (ambient = blue; sprayed = olive) or *E. cheiranthoides* (ambient = orange; sprayed = purple) plants in the 2023 field experiment. Points represent observations on individual plants, while lines and shaded show mean model predictions and 95% confidence intervals from negative binomial generalized additive models (GAMs). GAMs were fitted to count data after exclusion of weeks with zero observations. Each panel (a-h) shows treatment-specific predictions from the same model, which included insecticide as fixed effect. All models used ‘days in the field’ as temporal variable, but the x-axes show ‘week’ for easier comparison with other results. No model could be fit for crucifer shield bugs on insecticide-treated plants due to insufficient observations. See Supplementary Table S4 for test statistics of each model.

**Figure S5.**
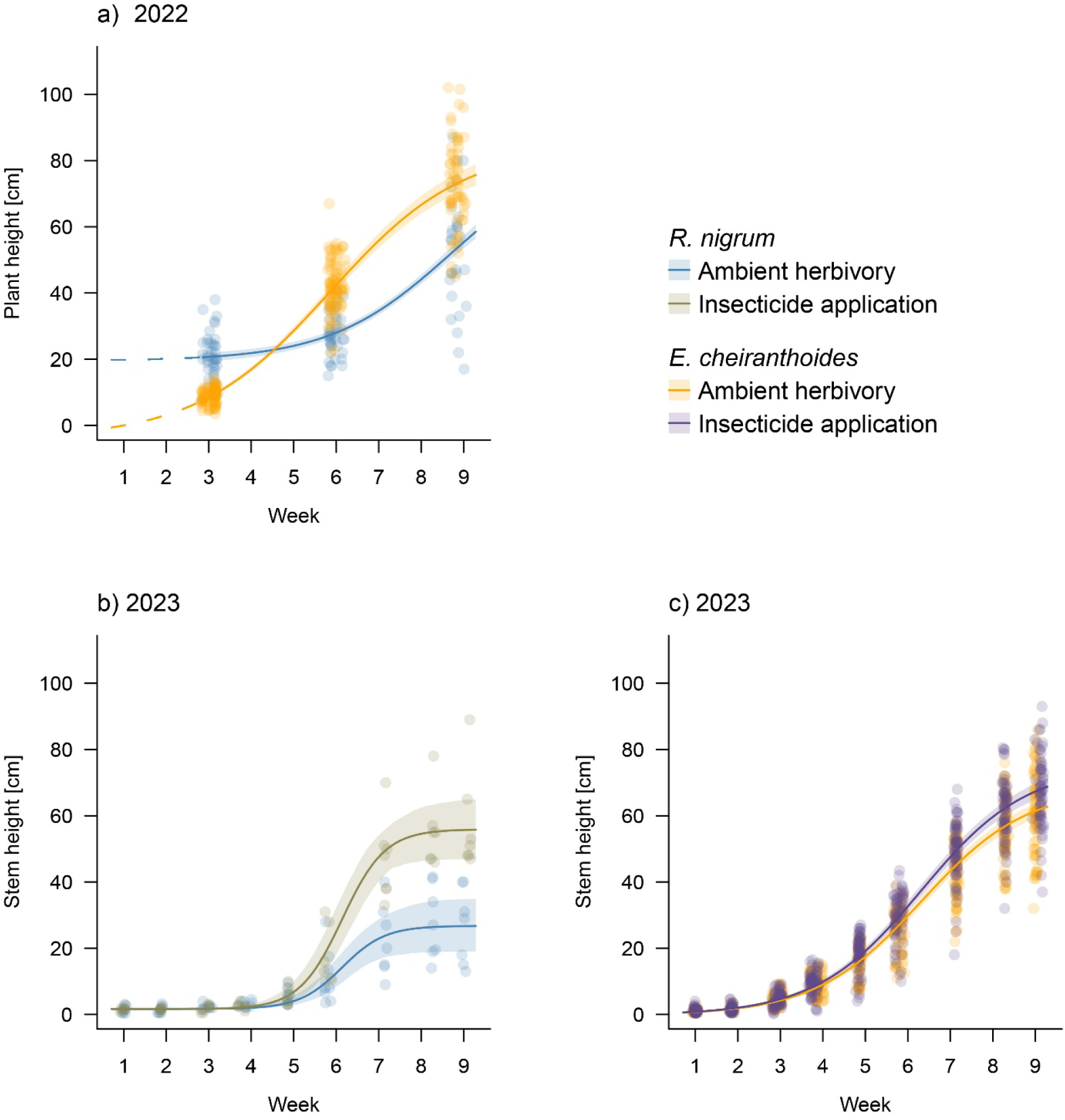
a) Plant height growth over nine weeks for *R. nigrum* and *E. cheiranthoides* in the 2022 field experiment. Points are individual plant measurements, and solid lines are model fits from a four-parameter logistic regression. Shaded areas indicate 95% confidence intervals (population prediction intervals). Plant height, measured from the root base to the tallest point of the plant, was first recorded in week 3; thus, the initial growth period (dashed lines) represents a backward extrapolation to the origin based on the fitted logistic function. b-c) Stem height growth for *R. nigrum* and *E. cheiranthoides* under ambient herbivory and insecticide treatment in the 2023 field experiment. Stem height was measured from the root base to the apical meristem of each plant, resulting in lower initial height values compared to 2022.

**Figure S6.**
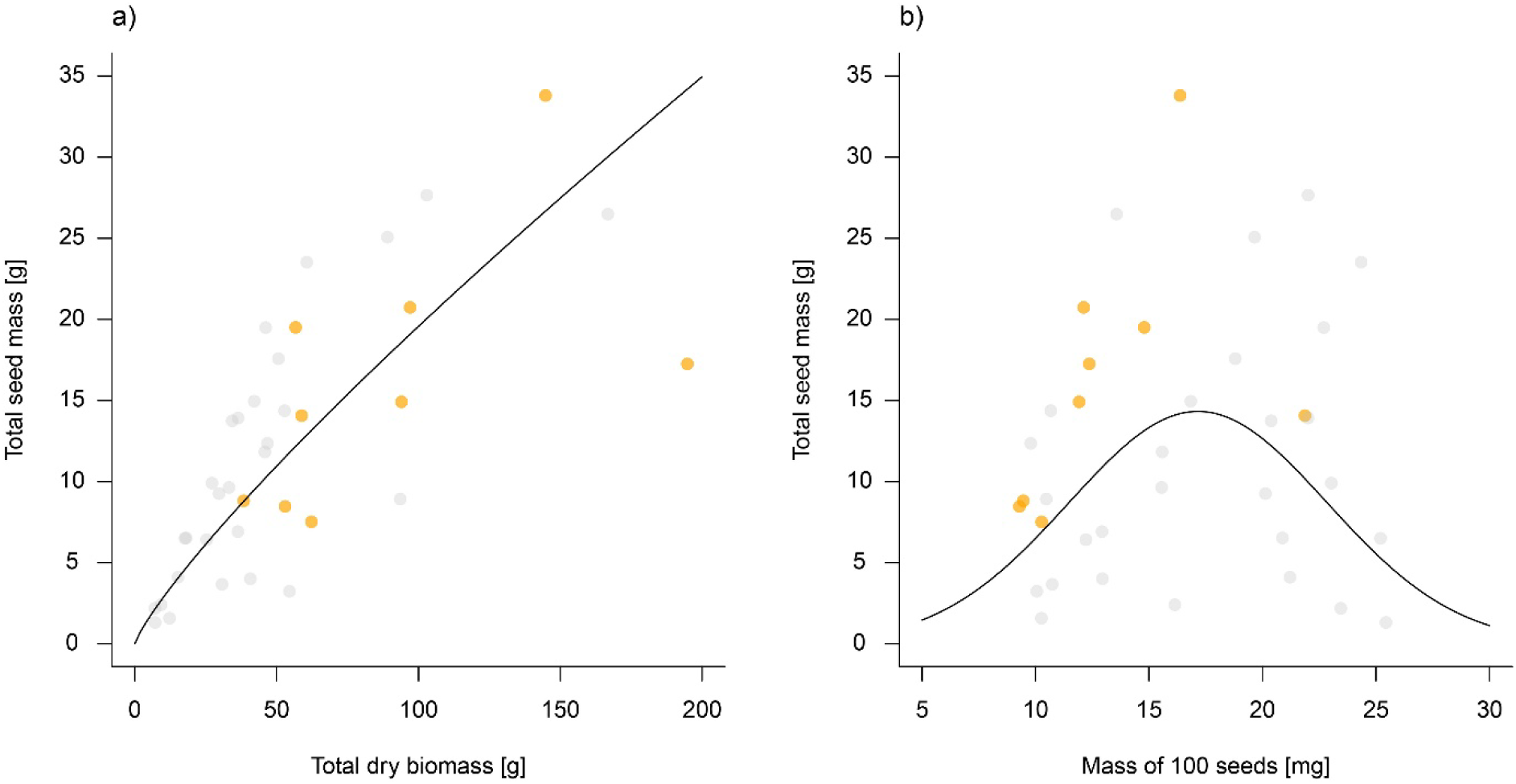
a) Relationship between total seed mass and aboveground plant biomass for *E. cheiranthoides* plants in the 2022 field experiment. Plants from block 7 were removed from the field intact and maintained in a greenhouse until seed ripening. Points represent individual plants, each corresponding to a different genotype. The solid line shows a linear fit between ln-transformed total seed mass and ln-transformed total biomass (F_1,36_ = 77.48, p<0.001, R^2^ = 0.67). Genotypes included in the main analyses are highlighted in orange, whereas excluded genotypes are shown grey. b) Relationship between total seed mass and individual seed mass. The solid line shows a quadratic fit between ln-transformed total seed mass and the mass of 100 seeds (quadratic term: F_1,35_ = 7.15, p=0.011, R^2^ = 0.12).

**Figure S7.**
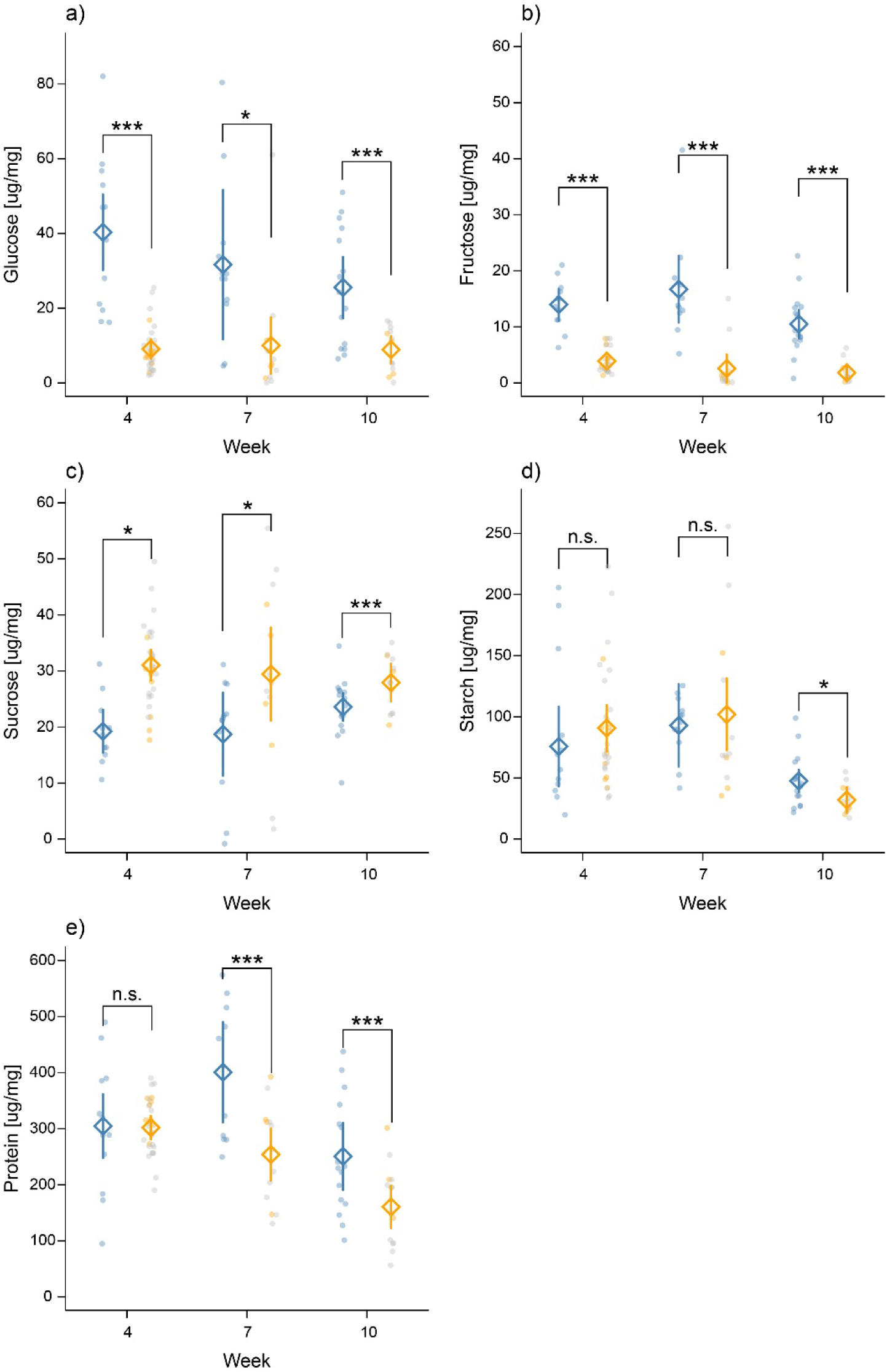
Concentrations of soluble sugars (a-c), starch (d), and total protein (d) in leaf tissue of *R. nigrum* (blue) and *E. cheiranthoides* (orange) harvested in weeks 4, 7, and 10 of the 2022 field experiment. Points are values for individual plants, while diamond symbols and lines are means and 95% confidence intervals from a linear model. Brackets highlight significant linear contrasts at the p < 0.05 level. For *E. cheiranthoides*, a random subset of plants from the full experiment was analysed, including genotypes excluded for other analyses in this study (grey points). Genotypes included in the main analyses are highlighted in orange, whereas excluded genotypes are shown grey.

**Figure S8.**
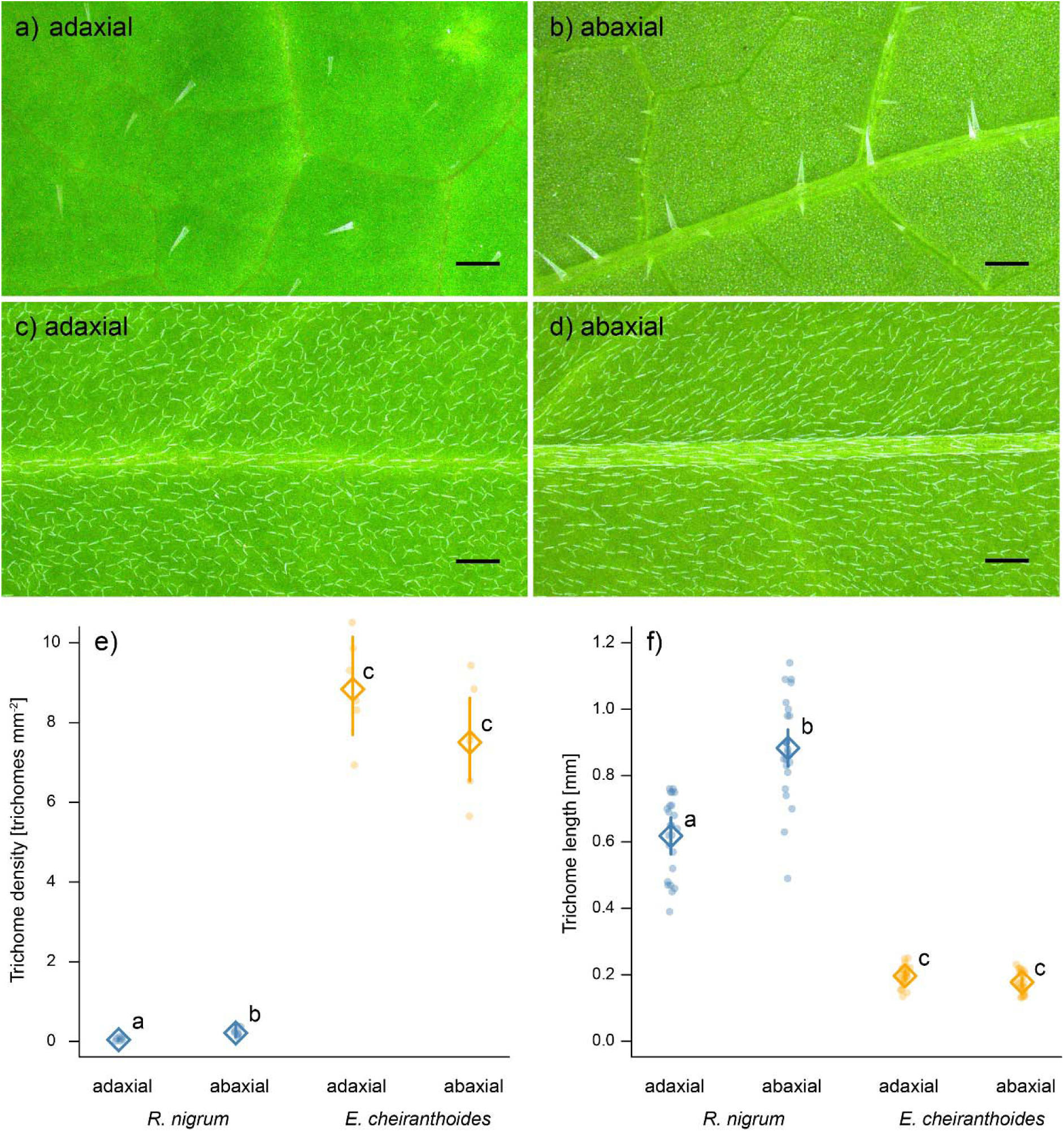
Photographs of the adaxial (upper) and abaxial (lower) leaf sides of 4-week-old *R. nigrum* (a-b) and *E. cheiranthoides* (c-d) plants growing under greenhouse conditions. Scale bars correspond to distances of 1 mm. Six photographs per side and species were used to quantify trichome densities (e) and trichome length (f). Points are values for individual plants, while diamonds and vertical lines are the means and 95% confidence intervals from linear models. Letters highlight significant linear contrasts at the p < 0.05 level.

**Figure S9.**
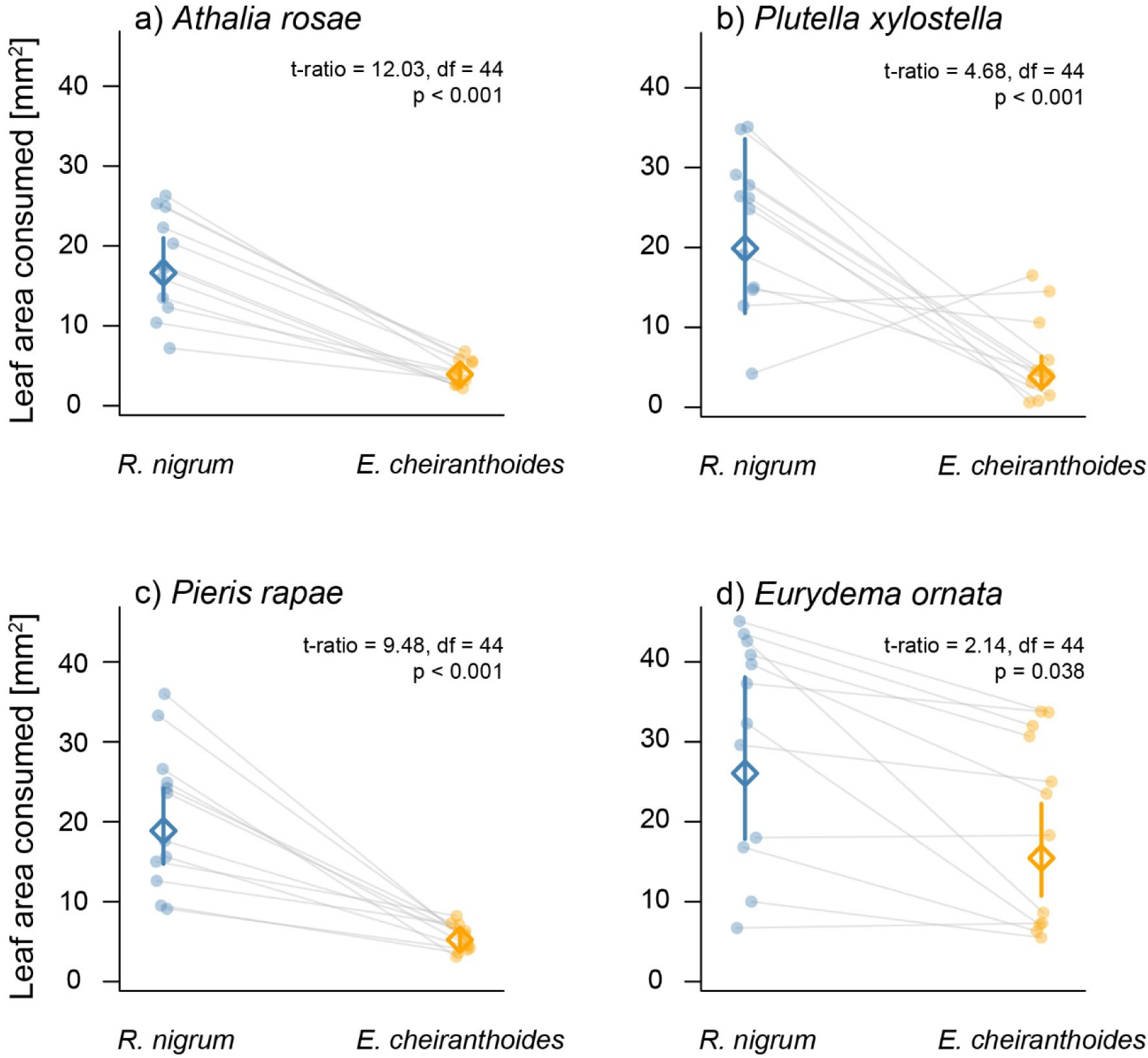
Relative preference of a) larvae of the turnip sawfly *Athalia rosae*, b) larvae of *P. xylostella*, c) larvae of the cabbage white butterfly *Pieris rapae*, or d), adult *E. ornata* in two-choice laboratory assays. Choice was quantified as leaf area eaten from pairs of leaf discs over 24 hours. Leaf discs (45 mm^2^ area) came from 4-week-old *R. nigrum* and *E. cheiranthoides* plants grown under greenhouse conditions and were offered to herbivores in Petri dishes. Points are damage areas for individual leaf discs, with grey lines connecting pairs of leaf discs from the same dish. Diamonds and vertical lines are the means and 95% confidence from a linear mixed-effects model, while statistical tests are linear contrasts comparing consumption of the two plant species for each herbivore.

## References

Agrawal, A. A., A. P. Hastings, M. T. J. Johnson, J. L. Maron, and J.-P. Salminen. 2012a. Insect herbivores drive real-time ecological and evolutionary change in plant populations. Science 338:113–116.

Agrawal, A. A., G. Petschenka, R. A. Bingham, M. G. Weber, and S. Rasmann. 2012b. Toxic cardenolides: chemical ecology and coevolution of specialized plant–herbivore interactions. New Phytologist 194:28–45.

Atsatt, P. R., and D. J. O’Dowd. 1976. Plant defense guilds. Science (New York, N.Y.) 193:24–29.

Barbosa, P., J. Hines, I. Kaplan, H. Martinson, A. Szczepaniec, and Z. Szendrei. 2009. Associational resistance and associational susceptibility: having right or wrong neighbors. Annual Review of Ecology, Evolution, and Systematics 40:1–20.

Bates, D., M. Mächler, B. Bolker, and S. Walker. 2015. Fitting linear mixed-effects models using lme4. Journal of Statistical Software 67.

Bekaert, M., P. P. Edger, C. M. Hudson, J. C. Pires, and G. C. Conant. 2012. Metabolic and evolutionary costs of herbivory defense: systems biology of glucosinolate synthesis. New Phytologist 196:596–605.

Bidart-Bouzat, M. G., and D. J. Kliebenstein. 2008. Differential levels of insect herbivory in the field associated with genotypic variation in glucosinolates in *Arabidopsis thaliana*. Journal of Chemical Ecology 34:1026–1037.

Clements, F. E., and G. W. Goldsmith. 1924. The phytometer method in ecology: the plant and community as instruments. The Carnegie Institution of Washington, Washington, D.C., USA.

Crawley, M. J. 1989. Insect herbivores and plant population dynamics. Annual Review of Entomology 34:531–564.

Dobler, S., S. Dalla, V. Wagschal, and A. A. Agrawal. 2012. Community-wide convergent evolution in insect adaptation to toxic cardenolides by substitutions in the Na,K-ATPase. Proceedings of the National Academy of Sciences 109:13040–13045.

Edger, P. P., H. M. Heidel-Fischer, M. Bekaert, J. Rota, G. Glöckner, A. E. Platts, D. G. Heckel, J. P. Der, E. K. Wafula, M. Tang, J. A. Hofberger, A. Smithson, J. C. Hall, M. Blanchette, T. E. Bureau, S. I. Wright, C. W. dePamphilis, M. Eric Schranz, M. S. Barker, G. C. Conant, N. Wahlberg, H. Vogel, J. C. Pires, and C. W. Wheat. 2015. The butterfly plant arms-race escalated by gene and genome duplications. Proceedings of the National Academy of Sciences 112:8362–8366.

Ehrlich, P. R., and P. H. Raven. 1964. Butterflies and plants: a study in coevolution. Evolution 18:586–608.

Fahey, J. W., A. T. Zalcmann, and P. Talalay. 2001. The chemical diversity and distribution of glucosinolates and isothiocyanates among plants. Phytochemistry 56:5–51.

Fox, J., and S. Weisberg. 2019. An R companion to applied regression. Third Edition. Sage, Thousand Oaks CA.

Gómez, J. M. 2005. Non-additive effects of herbivores and pollinators on *Erysimum mediohispanicum* (Cruciferae) fitness. Oecologia 143:412–418.

Hambäck, P. A., and A. P. Beckerman. 2003. Herbivory and plant resource competition: a review of two interacting interactions. Oikos 101:26–37.

Herms, D. A., and W. J. Mattson. 1992. The dilemma of plants: to grow or defend. The Quarterly Review of Biology 67:283.

Huang, X., J. A. A. Renwick, and K. Sachdev-Gupta. 1993. A chemical basis for differential acceptance of *Erysimum cheiranthoides* by two *Pieris* species. Journal of Chemical Ecology 19:195–210.

Jeffries, M. J., and J. H. Lawton. 1984. Enemy free space and the structure of ecological communities. Biological Journal of the Linnean Society 23:269–286.

Johnson, M. T. J. 2011. Evolutionary ecology of plant defences against herbivores: plant defences against herbivores. Functional Ecology 25:305–311.

Karasov, T. L., E. Chae, J. J. Herman, and J. Bergelson. 2017. Mechanisms to mitigate the trade-off between growth and defense. The Plant Cell 29:666–680.

Lenth, R. V. 2023. emmeans: Estimated Marginal Means, aka Least-Squares Means.

Levin, D. A. 1973. The role of trichomes in plant defense. The Quarterly Review of Biology 48:3–15.

Lüdecke, D., M. Ben-Shachar, I. Patil, P. Waggoner, and D. Makowski. 2021. performance: An R Package for Assessment, Comparison and Testing of Statistical Models. Journal of Open Source Software 6:3139.

Mauricio, R., and M. D. Rausher. 1997. Experimental manipulation of putative selective agents provides evidence for the role of natural enemies in the evolution of plant defense. Evolution 51:1435–1444.

Mertens, D., K. Bouwmeester, and E. H. Poelman. 2021. Intraspecific variation in plantDassociated herbivore communities is phylogenetically structured in Brassicaceae. Ecology Letters 24:2314–2327.

Myers, J. H., and R. M. Sarfraz. 2017. Impacts of insect herbivores on plant populations. Annual Review of Entomology 62:207–230.

Petschenka, G., R. Halitschke, T. Züst, A. Roth, S. Stiehler, L. Tenbusch, C. Hartwig, J. F. Moreno Gámez, R. Trusch, J. Deckert, K. Chalušová, A. Vilcinskas, and A. Exnerová. 2022. Sequestration of defenses against predators drives specialized host plant associations in preadapted milkweed bugs (Heteroptera: Lygaeinae). The American Naturalist 199:E211–E228.

Petschenka, G., V. Wagschal, M. Von Tschirnhaus, A. Donath, and S. Dobler. 2017. Convergently evolved toxic secondary metabolites in plants drive the parallel molecular evolution of insect resistance. The American Naturalist 190:S29–S43.

Piersanti, S., M. Rebora, L. Ederli, S. Pasqualini, and G. Salerno. 2020. Role of chemical cues in cabbage stink bug host plant selection. Journal of Insect Physiology 120:103994.

Polatschek, A. 2010. Revision der Gattung Erysimum (Cruciferae): Teil 1. Russland, die Nachfolgestaaten der USSR ( excl. Georgien, Armenien, Azerbaidzan), China, Indien, Pakistan, Japan und Korea. Annalen des Naturhistorischen Museums Wien B 111:181–275.

Polatschek, A. 2013. Revision der Gattung Erysimum (Cruciferae): Teil 5. Nord-, West-, Zentraleuropa, Rumänien und westliche Balkan-Halbinsel bis Albanien. Annalen des Naturhistorischen Museums Wien B 115:75–218.

R Core Team. 2023. R Statistical Software.

Ratzka, A., H. Vogel, D. J. Kliebenstein, T. Mitchell-Olds, and J. Kroymann. 2002. Disarming the mustard oil bomb. Proceedings of the National Academy of Sciences 99:11223–11228.

Sachdev-Gupta, K., C. D. Radke, J. A. A. Renwick, and M. B. Dimock. 1993. Cardenolides from *Erysimum cheiranthoides*: feeding deterrents to *Pieris rapae* larvae. Journal of Chemical Ecology 19:1355–1369.

Sato, Y., R. Shimizu-Inatsugi, M. Yamazaki, K. K. Shimizu, and A. J. Nagano. 2019. Plant trichomes and a single gene GLABRA1 contribute to insect community composition on field-grown Arabidopsis thaliana. BMC Plant Biology 19:1–12.

Scherber, C., P. N. Mwangi, V. M. Temperton, C. Roscher, J. Schumacher, B. Schmid, and W. W. Weisser. 2006. Effects of plant diversity on invertebrate herbivory in experimental grassland. Oecologia 147:489–500.

Simms, E. L., and M. D. Rausher. 1987. Costs and benefits of plant resistance to herbivory. The American Naturalist 130:570–581.

Speed, M. P., A. Fenton, M. G. Jones, G. D. Ruxton, and M. A. Brockhurst. 2015. Coevolution can explain defensive secondary metabolite diversity in plants. New Phytologist 208:1251–1263.

Sporer, T., J. Körnig, N. Wielsch, S. Gebauer-Jung, M. Reichelt, Y. Hupfer, and F. Beran. 2021. Hijacking the mustard-oil bomb: how a glucosinolate-sequestering flea beetle copes with plant myrosinases. Frontiers in Plant Science 12:645030.

Strauss, S. Y., J. A. Rudgers, J. A. Lau, and R. E. Irwin. 2002. Direct and ecological costs of resistance to herbivory. Trends in Ecology & Evolution 17:278–285.

Wang, K., Q. Rusman, E. van Bergen, and T. Züst. 2025. Effects of dual chemical defences on herbivory: Leaf damage and herbivore abundance in Erysimum cheiranthoides are negatively associated with cardenolides, but not glucosinolates.

Wang, K., and T. Züst. 2025. WithinDplant variation in chemical defence of Erysimum cheiranthoides does not explain Plutella xylostella feeding preference. Plant Biology:plb.13777.

War, A. R., M. G. Paulraj, T. Ahmad, A. A. Buhroo, B. Hussain, S. Ignacimuthu, and H. C. Sharma. 2012. Mechanisms of plant defense against insect herbivores. Plant Signaling & Behavior 7:1306–1320.

Wittstock, U., N. Agerbirk, E. J. Stauber, C. E. Olsen, M. Hippler, T. Mitchell-Olds, J. Gershenzon, and H. Vogel. 2004. Successful herbivore attack due to metabolic diversion of a plant chemical defense. Proceedings of the National Academy of Sciences of the United States of America 101:4859–4864.

Wood, S. N., N. Pya, and B. Säfken. 2016. Smoothing parameter and model selection for general smooth models. Journal of the American Statistical Association 111:1548–1563.

Younkin, G. C., M. L. Alani, A. PáezDCapador, H. D. Fischer, M. Mirzaei, A. P. Hastings, A. A. Agrawal, and G. Jander. 2024. Cardiac glycosides protect wormseed wallflower (*Erysimum cheiranthoides*) against some, but not all, glucosinolateDadapted herbivores. New Phytologist 242:2719–2733.

Zalucki, J. M., D. G. Heckel, P. Wang, S. Kuwar, D. G. Vassão, L. Perkins, and M. P. Zalucki. 2021. A generalist feeding on Brassicaceae: it does not get any better with selection. Plants 10:954.

Züst, T., and A. A. Agrawal. 2017. Trade-offs between plant growth and defense against insect herbivory: an emerging mechanistic synthesis. Annual Review of Plant Biology 68:513–534.

Züst, T., C. Heichinger, U. Grossniklaus, R. Harrington, D. Kliebenstein, and L. Turnbull. 2012. Natural enemies drive geographic variation in plant defenses. Science 338:116–119.

Züst, T., B. Joseph, K. K. Shimizu, D. J. Kliebenstein, and L. A. Turnbull. 2011. Using knockout mutants to reveal the growth costs of defensive traits. Proceedings of the Royal Society B: Biological Sciences 278:2598–2603.

Züst, T., S. Rasmann, and A. A. Agrawal. 2015. Growth-defense tradeoffs for two major anti-herbivore traits of the common milkweed *Asclepias syriaca*. Oikos 124:1404–1415.

Züst, T., S. R. Strickler, A. F. Powell, M. E. Mabry, H. An, M. Mirzaei, T. York, C. K. Holland, P. Kumar, M. Erb, G. Petschenka, J.-M. Gómez, F. Perfectti, C. Müller, J. C. Pires, L. A. Mueller, and G. Jander. 2020. Independent evolution of ancestral and novel defenses in a genus of toxic plants (*Erysimum*, Brassicaceae). eLife 9:e51712.

## Supplementary References

Aliabadi, A., Renwick, J. A. A., Whitman, D. W. (2002). Sequestration of glucosinolates by harlequin bug *Murgantia histrionica*. Journal of Chemical Ecology 28, 1749–1762.

Bolker, B. M. (2008). Ecological Models and Data in R. Princeton University Press.

Brooks, M., E., Kristensen, K., Benthem, K., J. ,van, Magnusson, A., Berg, C., W., Nielsen, A., Skaug, H., J., Mächler, M., & Bolker, B., M. (2017). glmmTMB balances speed and flexibility among packages for zero-inflated generalized linear mixed modeling. The R Journal, 9(2), 378. 10.32614/RJ-2017-066

Clissold, F. J., Sanson, G. D., & Read, J. (2006). The paradoxical effects of nutrient ratios and supply rates on an outbreaking insect herbivore, the Australian plague locust. Journal of Animal Ecology 75, 1000–1013.

Friedman, J., Hastie, T., & Tibshirani, R. (2010). Regularization Paths for Generalized Linear Models via Coordinate Descent. Journal of Statistical Software 33, 1–22.

Machado, R. A. R., Ferrieri, A. P., Robert, C. A. M., Glauser, G., Kallenbach, M., Baldwin, I. T., & Erb, M. (2013). Leaf-herbivore attack reduces carbon reserves and regrowth from the roots via jasmonate and auxin signaling. New Phytologist 200, 1234–1246.

Müller, C. & Wittstock, U. (2005). Uptake and turn-over of glucosinolates sequestered in the sawfly *Athalia rosae*. Insect Biochemistry and Molecular Biology 35, 1189–1198.

Oksanen, J., Simpson, G. L., Blanchet, F. G., Kindt, R., Legendre, P., Minchin, P., O’Hara, R. B., Solymos, P., Stevens, M. H. H., Szoecs, E., Wagner, H., Barbour, M., Bedward, M., Bolker, B., Borcard, D., Borman, T., Carvalho, G., Chirico, M., De Caceres, M., … Weedon, J. (2025). vegan: Community ecology package [Computer software].

Schneider C.A., Rasband W.S., Eliceiri K.W. (2012). NIH Image to ImageJ: 25 years of image analysis. Nature Methods 9, 671–675.

Smith, A. M., & Zeeman, S. C. (2006). Quantification of starch in plant tissues. Nature Protocols 1, 1342–1345.

Velterop, J. S., & Vos, F. (2001). A rapid and inexpensive microplate assay for the enzymatic determination of glucose, fructose, sucrose, L-malate and citrate in tomato (*Lycopersicon esculentum*) extracts and in orange juice. Phytochemical Analysis 12, 299–304.

Venables, W. N., & Ripley, B. D. (2002). Modern applied statistics with S. Fourth Edition. Springer, New York.

